# Rationally Engineered Coronavirus-mimicking Protein Nanocage Platform for Glioblastoma Targeted Delivery and Immunity Regulation

**DOI:** 10.1101/2025.08.08.669313

**Authors:** Jiaqi Li, Jian Feng, Yue Ren, Liping Wang, Jie Wang, Xing Liu, Shuoqi Tian, Xinyi Yuan, Jiaoyan Li, Jinxia Huang, Chunhui Liu, Yitian Du, Yan Xia, Shuang Jia, Yinan Sun, Shubin Li, Rile Wu, Liyao Wang, Xinyu Li

## Abstract

Glioblastoma (GBM) is one of the most aggressive and treatment-resistant brain tumors, owing to the dual challenges of the blood-brain barrier (BBB) and a profoundly immunosuppressive tumor microenvironment. However, few existing strategies are capable of simultaneously overcoming these two barriers, underscoring a critical scientific gap and the need for innovative therapeutic platforms that enable both effective BBB traversal and tumor immune reprogramming. Here, we present a coronavirus-mimicking protein nanocage platform (EcomPC) derived from *Thermotoga maritima* encapsulin, rationally engineered for the targeted and sustained delivery of interferon-α (IFN-α) to GBM lesions. Through strategic insertion of flexible linkers, cysteine-to-serine mutations, and modular surface functionalization via SpyTag/SpyCatcher chemistry, EcomPC enables intracellular self-assembly of IFN-α and glioma-specific targeting. In orthotopic GBM models, EcomPC–IFN demonstrates efficient BBB translocation, selective tumor accumulation, and potent anti-tumor efficacy. Mechanistically, localized IFN-α release induces tumor cell apoptosis and reprograms the immune microenvironment—characterized by increased CD8⁺ T cell infiltration, decreased Foxp3⁺ regulatory T cells, and a favorable chemokine shift. These therapeutic effects are not recapitulated by free IFN-α or untargeted nanocages, underscoring the essential role of both structural mimicry and ligand-guided delivery. Collectively, this work establishes EcomPC as a programmable, virus-inspired protein delivery platform capable of overcoming key physiological barriers in brain tumor immunotherapy, and lays the foundation for its broader application in CNS-targeted biologic delivery.

## Introduction

Glioblastoma (GBM) is the most aggressive form of primary brain cancer, accounting for approximately 45% of intracranial tumors, with an incidence rate of 30 to 100 cases per million^1–4^. The median survival time for patients with GBM is approximately 15 months, and the overall 5-year survival rate is less than 10%, underscoring the significant impact of this disease on public health^5,6^. Current treatment strategies typically involve a combination of surgery and adjuvant therapies, such as radiation and chemotherapy^2,3,7^. Further combination of therapeutic proteins and anti-bodies with small molecular drugs could significantly improve the anti-GBM effects^8,9^. However, the therapeutic efficacy of these treatments is often hindered by challenges related to drug accumulation and internalization within tumor lesions, primarily due to the presence of the blood-brain barrier (BBB) and the blood-brain tumor barrier (BBTB)^6,10,11^. Consequently, nearly all macromolecules and approximately 98% of small molecule drugs are unable to effectively penetrate GBM lesions^6^.

Although various targeted drug delivery systems, including liposomes^12^, polymeric nanoparticles^13^, and antibody-drug conjugates^14^, have been developed to enhance drug access to GBM, few have achieved meaningful clinical outcomes^4^. One of the primary reasons for this therapeutic failure is the profoundly immunosuppressive nature of the GBM microenvironment^1,15^. Suppressed innate and adaptive immune responses allow tumor cells to evade immunosurveillance and resist immunotherapeutic interventions^1,3,15^. Therefore, next-generation GBM therapies must address two critical bottlenecks: efficient penetration across the BBB and reprogramming of the local immune landscape to overcome immunosuppression. Intriguingly, certain neurotropic viruses, most notably coronaviruses, have evolved mechanisms to breach the BBB and establish infections within neural tissues^16–18^. Their multivalent, spike-decorated surfaces engage specific host receptors and exploit vesicular transport pathways, facilitating transcytosis through endothelial layers^16,19^. These features offer a powerful blueprint for designing virus-inspired nanocarriers capable of mimicking viral tropism for CNS-targeted drug delivery^20^. Moreover, coronaviruses are known to stimulate potent innate and adaptive immune responses through pattern recognition receptors (PRRs)^21,22^ and T cell-mediated immunity^23^, suggesting that their structural features can intrinsically modulate host immunity^20^.

Inspired by this biological paradigm, we hypothesized that rationally engineered protein nanocages, incorporating virus-mimicking surface topologies^19^ and tumor-targeting ligands^6^, could recapitulate key aspects of viral transport and immunogenicity. Unlike synthetic nanoparticles, protein-based nanocages offer genetically encoded precision, uniform architecture, high biocompatibility, and modularity—allowing for tight control over particle size, surface presentation, and cargo loading^24–26^. These properties are essential not only for efficient navigation across the BBB but also for tuning interactions within the tumor immune microenvironment.

To test this hypothesis, we engineered a coronavirus-inspired protein delivery platform, termed EcomPC (**E**ncapsulin-based **C**or**o**navirus-**M**imicking **P**rotein **C**age). Derived from the self-assembling encapsulin scaffold^27–30^, EcomPC was structurally reconfigured to incorporate flexible linkers and eliminate disulfide constraints, thereby increasing its cargo capacity and structural adaptability^31^. Internally, therapeutic proteins were loaded using intein-mediated self-assembly; externally, cell-targeting peptides were covalently displayed via SpyTag/SpyCatcher chemistry to generate spike-like topologies^32^. This dual-level engineering mimics both the cargo-packaging efficiency and targeting precision of natural viruses.

To this end, we developed EcomPC, a coronavirus-mimicking, encapsulin-derived protein cage platform, engineered for targeted, sustained delivery of bioactive payloads such as interferon-α (IFN-α) to GBM lesions. By combining intracellular protein encapsulation via split-intein splicing with surface functionalization through SpyTag/SpyCatcher chemistry, EcomPC offers a programmable, biomimetic nanoplatform for overcoming physiological barriers and reshaping tumor immunity.

## Results

### Design and characteristics of EcomPC

To construct the EcomPC platform, we selected the encapsulin protein from *Thermotoga maritima* as a structural scaffold due to its well-characterized ability to self-assemble into hollow icosahedral nanocages composed of 60 identical subunits^28,29,27,28,33^. Structural analysis revealed that residues Phe30, Leu34, Leu233, and Ile249 form a conserved hydrophobic core that is likely critical for maintaining cage stability and guiding correct assembly^34^. Mutations at these positions abolished nanocage formation (Fig. S2), underscoring their essential structural role. In contrast, a double point mutation (C130S/C196S) located in the P-domain and A-domain, respectively, preserved cage formation while reducing disulfide-mediated crosslinking. Additionally, insertion of a rationally designed flexible linker between residues P41 and Y42 (within the E-loop) further enhanced conformational adaptability without compromising structural integrity (Fig.1D, Fig. S1 and Fig. S2) ^31^. Dynamic light scattering (DLS) and transmission electron microscopy (TEM) confirmed successful cage assembly, revealing an average diameter of 36.38 ± 19.63 nm, significantly larger than the ∼20–25 nm diameter of wild-type encapsulin^27,33^, indicating an expanded internal cavity and enhanced structural plasticity (Fig. 1 E, I and Table 1).

**Fig 1.**
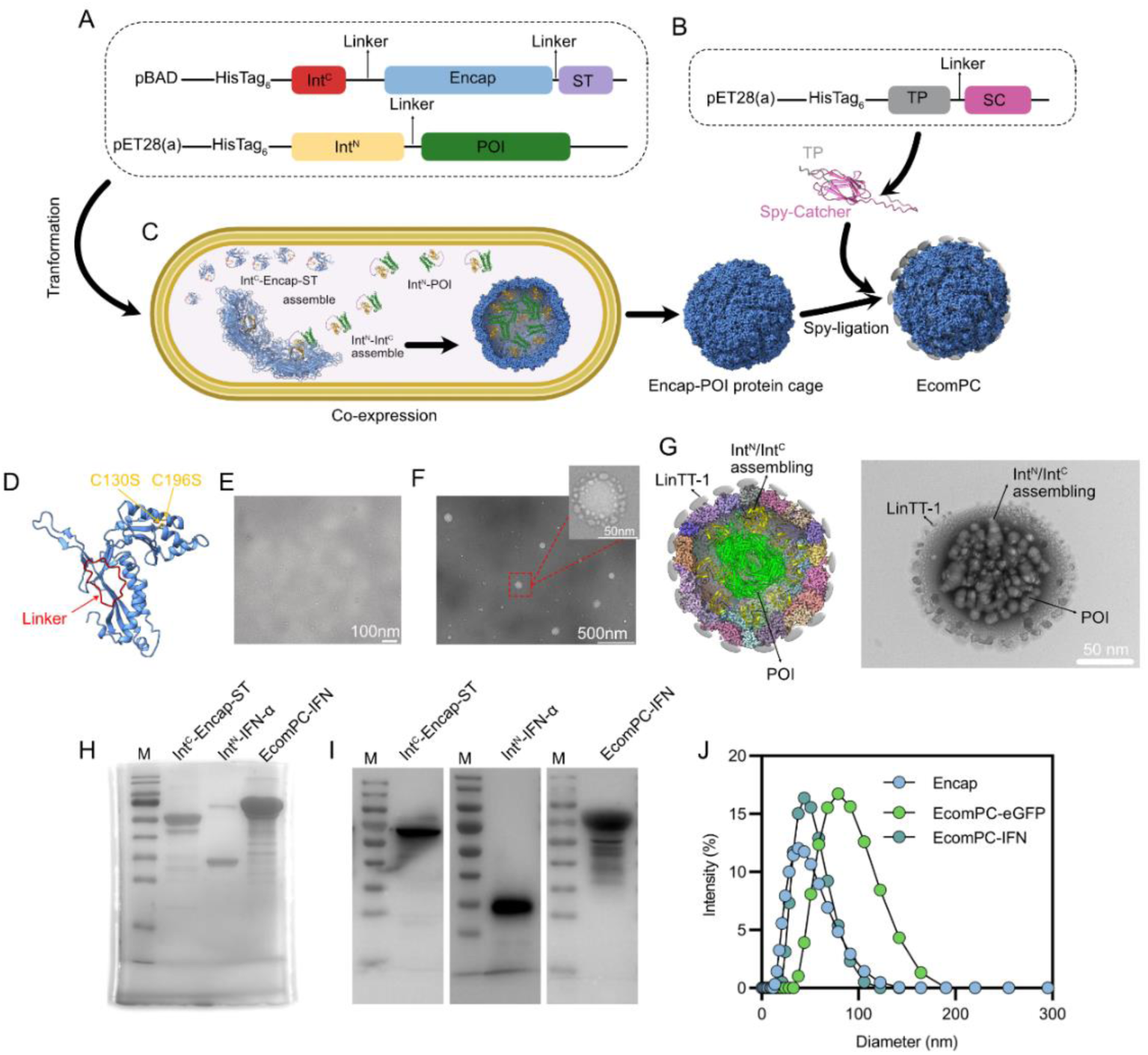
Design, assembly, and characterization of the virus-mimicking EcomPC nanocage platform. A. Schematic illustration of the intein-mediated cargo encapsulation strategy. The POI was fused to the C-terminal of Int^N^, while the encapsulin scaffold was fused to the C-terminal of Int^C^ and a C-terminal SpyTag for downstream surface functionalization. Co-expression in *E. coli* BL21(DE3) enabled intracellular protein cage assembly and spontaneous intein-mediated ligation of cargo POI. B. Schematic of EcomPC exterior functionalization. SpyCatcher-tagged targeting peptides (SC-TP) were conjugated to the SpyTag-presenting cage surface, enabling modular and stable attachment of ligands. C. Overview of the modular EcomPC assembly strategy combining internal protein encapsulation with external ligand conjugation. D. Encap monomer structure predicted by Alphafold 3. A flexible linker was inserted between residues P41 and Y42, and internal cysteines C130 and C196 were mutated to serines to reduce disulfide crosslinking and enhance conformational flexibility. E. TEM image of natural Encap protein cage. F. TEM of EcomPC showing uniform spherical particles with corona-like surface projections reminiscent of enveloped viruses. G. High-voltage TEM imaging revealing dense internal cargo distribution within disrupted cages, confirming successful encapsulation of POI. H. SDS-PAGE results showing the co-expression of Int^C^-Encap-ST and Int^N^–IFN, resulting in a fused ∼88–90 kDa band upon successful splicing and cage assembly. I. Western bolt confirms the co-expression of Int^C^-Encap-ST and Int^N^–IFN. J. DLS analysis of particle size distributions. Cargo loading altered cage dimensions: IFN-α-loaded EcomPC measured ∼43.49 ± 16.15 nm, and eGFP-loaded cages reached ∼75.50 ± 26.83 nm, demonstrating cargo-dependent size tunability.

**Table 1.** Hydrodynamic diameter and zeta potential of engineered Encap, EcomPC-IFN and EcomPC-eGFP.

|  | Z-Average (d.nm) | St Dev (d.nm) | Pdl | Zeta potenti (mv) | Conductivity (ms/cm) | St Dev(mv) |
| --- | --- | --- | --- | --- | --- | --- |
| Encap | 36.38 | 19.63 | 0.155 | -6.7 | 23.3 | 5.32 |
| EcomPC-eGFP | 75.50 | 26.83 | 0.080 | -23.6 | 0.0469 | 5.78 |
| EcomPC-IFN-α | 43.49 | 16.15 | 0.097 | -13.1 | 2.48 | 3.34 |

Intein-mediated protein splicing system could medium protein assemble during expression without disrupted protein functions^35^. Therefore, to enable selective cargo encapsulation, the protein of interest (POI) was fused with a split intein (Int^N^) and the corresponding C-terminal intein fragment (Int^C^) was genetically fused to the N-terminus of encapsulin via a flexible linker (Fig. 1A). The POI-Int^N^ construct and the Int^C^-Encap-ST construct was co-expressed in *E. coli* BL21(DE3) enabled intracellular assembly and spontaneous trans-splicing between Int^N^ and Int^C35^, resulting in efficient encapsulation of the POI within the protein nanocage (Fig. 1A, C). SDS-PAGE and Western blot analyses confirmed successful cage assembly, as co-expression of Int^C^–Encap–ST (55 kDa) and Int^N^– IFN (35 kDa) yielded a fusion product of ∼88–90 kDa, consistent with efficient intein-mediated splicing and encapsulation (Fig. 1H, I) ^27,36^.

The other reasons we choose Encap of *Thermotoga maritima* as a structural scaffold due to C-terminal targeting peptides (TPs) modification are well documented in this protein without disrupted protein assemble^28^. For exterior TPs functionalization, Spy-chemistry was selected rather than directly fusing TPs to C-terminal of Encap that because hard to mimic architecture of coronaviruses and endow cell-targeting capability in directly fusion paradigm^27,32^. Therefore, a SpyCatcher–targeting peptide (SC–TP) fusion was expressed in *E. coli*, purified, and subsequently incubated with POI-loaded cages. This facilitated isopeptide linkage via SpyTag/SpyCatcher chemistry, generating the fully assembled EcomPC construct (Fig. 1B, C and Fig. S3).

Coronavirus charactistics with spike proteins and hollow spherical structures^16,23^. The designed EcomPC in this study, the structural morphology of EcomPC was visualized and confirmed by TEM. The EcomPC exhibited uniform, spherical architecture with distinct spike-like surface protrusions, closely resembling the coronaviruses-like structure of enveloped viruses (Fig. 1F). High-voltage electron beams were applied to induce partial cage disruption to assess internal cargo distribution. Cross-sectional TEM imaging revealed densely packed interiors, confirming successful encapsulation of cargo proteins (Fig. 1G). Further DLS measurements revealed distinct size profiles depending on the encapsulated cargo protein. The average diameters of EcomPC loaded with IFN-α and eGFP were 43.49 ± 16.15 nm and 75.50 ± 26.83 nm, respectively, compared to 36.38 ± 19.63 nm for the empty encapsulin cage (Fig. 1J, Table 1). These results indicate that the overall particle size of EcomPC is tunable and correlates with the molecular weight and volume of the encapsulated proteins.

### In Vitro Evaluation of BBB Penetration and Cellular Uptake of EcomPC

As previously reported, both the receptor-binding affinity of coronavirus spike proteins^16^ and their overall structural organization^22^ play critical roles in facilitating BBB translocation and cellular uptake. We therefore hypothesized that the coronavirus-inspired structural architecture of EcomPC, combined with the multivalent display a GBM targeted and penetrating peptides, would synergistically promote BBB transcytosis and tumor-specific uptake. To assess the ability of EcomPC to traverse the BBB, we genetically encapsulated eGFP within the engineered nanocage and functionalized its surface with LinTT1, a glioblastoma-homing peptide identified via T7 phage display against recombinant human p32 (gC1qR)^6,37^. We further designed three control constructs: (i) native encapsulin loaded with eGFP (Encap-eGFP); (ii) EcomPC carrying a LinTT1 loss-of-function mutant (EcomPC_mut_- eGFP) (Fig. S5 and Table. S1); and (iii) a recombinant LinTT1-eGFP fusion protein lacking the nanocage framework (Fig. 2A).

**Fig 2.**
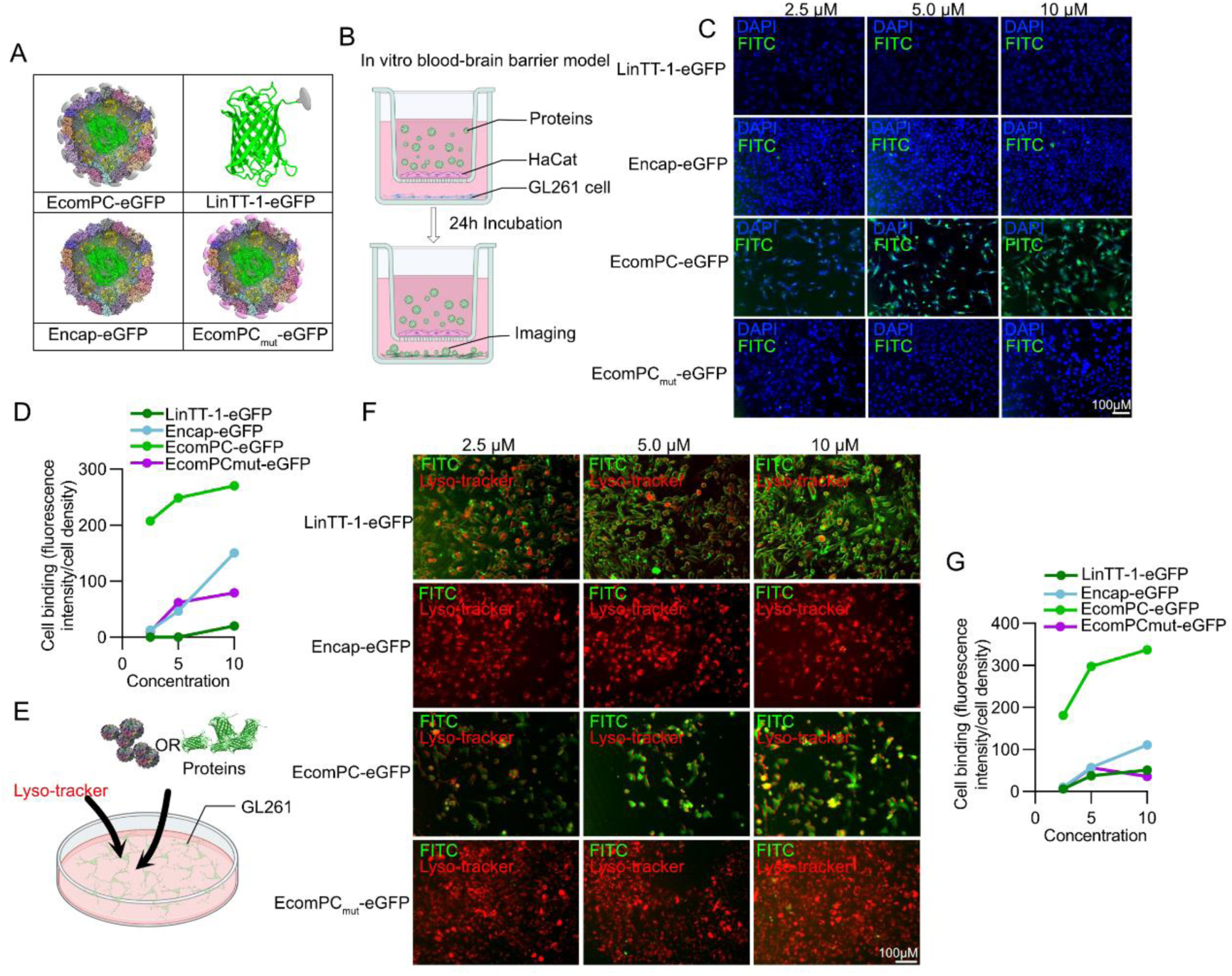
In vitro BBB and cell membrane permeable of EcomPC. A. Schematic illustration of engineered protein constructs. EcomPC-eGFP served as the experimental group, featuring encapsulated eGFP and surface LinTT-1 targeting. Three control constructs were used: LinTT1-eGFP, Encap-eGFP, and EcomPC_mut_-eGFP. B. Diagram of the in vitro transwell BBB model. Human HaCaT keratinocytes were cultured on the upper membrane insert, and GL261 glioma cells were seeded in the lower chamber. C. Representative fluorescence images of GL261 cells 24 h after treatment with the indicated constructs. D. Quantification of fluorescence intensity normalized to cell density, showing that EcomPC-eGFP achieved markedly enhanced translocation across the BBB model. E. Schematic of the cell membrane permeability assay. Constructs were incubated with GL261 cells for 6 h, followed by lysosomal staining with LysoTracker. F. Fluorescence microscopy images showing subcellular localization of each construct in GL261 cells. G. Quantitative analysis of intracellular fluorescence intensity per cell, confirming efficient cellular uptake.

We first examined cell viability following treatment to assess potential cytotoxicity of the experimental constructs. The results indicated that neither EcomPC-GFP nor the control groups induced detectable cytotoxic effects (Fig. S5). Based on this safety profile, we next evaluated the BBB permeability using a transwell co-culture model (Fig. 2B) ^37^. After EcomPC-eGFP administration and 24h incubation, GL261 cells of lower chamber exhibited robust intracellular fluorescence, indicating efficient transport across the in vitro BBB model (Fig. 2C, D). In contrast, fluorescence was minimal in groups treated with Encap-eGFP, EcomPC_mut_-eGFP, or LinTT1-eGFP fusion protein, highlighting the requirement of both the intact nanocage structure and functional LinTT1 ligand for effective BBB crossing (Fig. 2C, D).

Cell membrane permeability is a hallmark of coronaviruses^17,19^ and a critical feature for the effective function of VLP delivery system^20,38,39^. We therefore investigated whether the EcomPC platform possesses similar capabilities (Fig. 2E). Interestingly, unlike native coronaviruses, whose membrane-permeating ability is conferred by their spike proteins^19^, the LinTT1-modified eGFP construct, lacking a virus-like architecture, exhibited strong membrane association but failed to penetrate GL261 cells. This was evidenced by peripheral membrane fluorescence that intensified with increasing concentrations, yet remained restricted to the cell surface (Fig. 2F, G). Neither Encap–eGFP nor EcomPC_mut_–eGFP triggered appreciable fluorescence at any concentration, confirming the necessity of both the intact nanocage and functional LinTT1 ligand for uptake (Fig. 2F, G). In contrast, EcomPC– eGFP induced dose-dependent, diffuse cytoplasmic fluorescence, indicating successful membrane translocation and intracellular delivery (Fig. 2F, G). Moreover, unlike conventional antibody-based carriers that undergo endocytosis followed by lysosomal trafficking^20^, EcomPC demonstrated minimal lysosomal colocalization, suggesting efficient endosomal escape. This escape is critical to protect therapeutic proteins from degradation and ensure their bioactivity within the cytoplasm^20^. Taken together, these findings establish that ECoMPC’s virus-inspired geometry, coupled with targeted ligand presentation, enables not only BBB transcytosis but also active membrane penetration and cytoplasmic release, features essential for intracellular delivery of macromolecular therapeutics in CNS malignancies^20^.

### EcomPC Encapsulation Enhances the Anti-Tumor Efficacy of IFN-α in Glioma Cells

As an FDA-approved cytokine, IFN-α exerts broad anti-tumor effects, including inhibition of cell proliferation, induction of apoptosis, suppression of angiogenesis, and modulation of the immune response at the tumor site^40,41^. Despite these attributes, the clinical application of IFN-α for glioblastoma (GBM) remains severely limited by its poor penetration across the BBB, rapid systemic clearance, and dose-limiting toxicity^14,41^. Overcoming these challenges requires delivery systems that not only improve pharmacokinetics but also preserve bioactivity and enable precise targeting within the central nervous system^14,41^. Building upon our findings, we replaced eGFP with IFN-α to evaluate the therapeutic efficacy of EcomPC as a cytokine delivery platform. The anti-proliferative activity of EcomPC-encapsulated IFN-α was assessed in GL261 glioma cells and compared to native IFN-α, empty nanocages (Encap), and PBS. As expected, PBS and Encap showed negligible cytotoxicity, even at concentrations up to 20 μM (Fig. 3A and Fig. S6). Free IFN-α reduced cell viability by ∼50% at 20 μM, consistent with its known anti-tumor activity^42^. Notably, EcomPC-IFN achieved a similar cytotoxic effect at just 10 μM, demonstrating superior potency through nanocage-mediated delivery (Fig. 3A). Live/dead staining further revealed that EcomPC-IFN induced significant cell death at concentrations as low as 5 μM, whereas free IFN-α exhibited weaker cytotoxicity (Fig. 3B and Fig. S7). Flow cytometry analysis corroborated these results, showing a markedly higher proportion of dead cells in the EcomPC-IFN group compared to all controls (Fig. 3C). Importantly, this effect was tumor-selective: EcomPC-IFN did not induce same cytotoxicity in non-target HeLa cells at equivalent concentrations (Fig. S9, S10), highlighting the role of ligand-guided targeting and suggesting reduced off-target toxicity.

**Fig 3.**
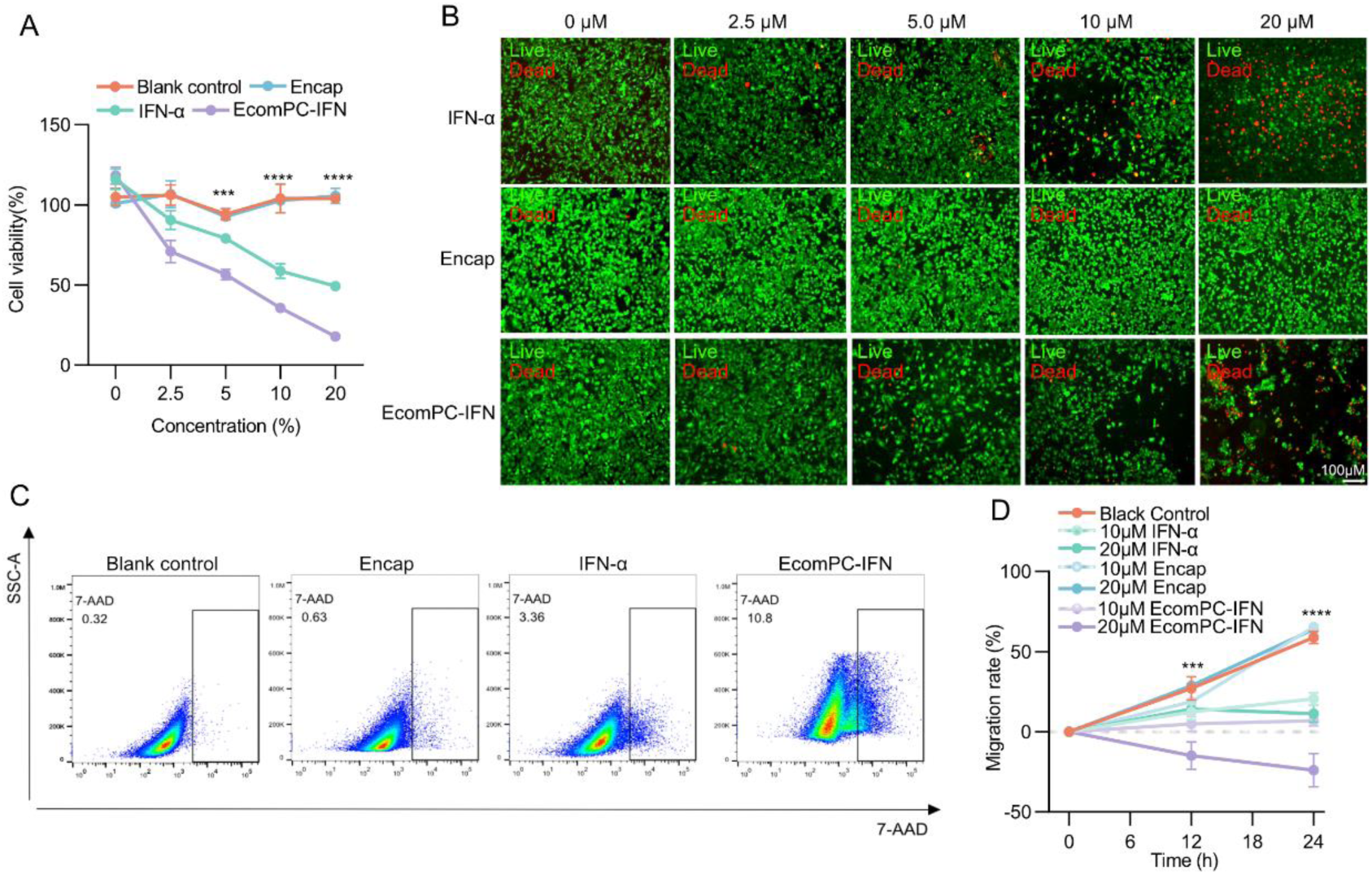
In vitro anti-tumor effects of EcomPC-IFN on GL261 glioma cells. A. Quantification of cell viability by CCK-8 assay after treatment with increasing concentrations (0-20 μM) of EcomPC-IFN, free IFN-α, Encap, or PBS. B. Representative live/dead staining images of GL261 cells following 6-hour treatment with the indicated formulations to assess cytotoxicity. C. Flow cytometry-based quantification of live and dead GL261 cells after treatment with EcomPC-IFN and control groups, confirming enhanced cytotoxicity. D. Quantitative analysis of GL261 cell migration assessed by wound-healing (scratch) assay after 24-hour treatment with EcomPC-IFN and controls; negative values indicate wound expansion due to cell death.

Glioblastoma (GBM) is the most aggressive primary brain tumor, characterized by rapid invasion and resistance to therapy^2^. Inhibiting tumor cell migration is therefore a critical therapeutic objective in GBM treatment^43^. We further conducted wound-healing assays to investigate the impact of EcomPC-IFN on glioma cell migration^44^. In PBS- and Encap-treated controls (10 and 20 μM), wound closure reached ∼73% after 24 hours, reflecting uninhibited migration (Fig. 3D). Free IFN-α treatment (10 or 20 μM) significantly impaired migration, reducing closure to <10% (Fig. 3D and Fig. S6). Notably, at 20 μM, EcomPC-IFN not only abrogated migration but induced pronounced cytotoxicity at the wound margins, yielding a negative migration index of –28% (Fig. 3D and Fig. S8) compare with other mutations (Fig. S8). Together, these findings demonstrate that EcomPC-IFN robustly enhances the anti-tumor efficacy of IFN-α in vitro, exerting synergistic effects on both cytotoxicity and cell motility. The nanocage-based delivery system not only potentiates the therapeutic function of IFN-α but also amplifies its anti-migratory effects, key features for combating the invasive nature of GBM.

### EcomPC-IFN BBB Permeability, Tumor Targeting, and Anti-GBM Activity in Orthotopic Mouse Model

To further evaluate the therapeutic potential of EcomPC-IFN in vivo, we examined its capacity to cross the BBB, selectively target GBM lesions, and elicit anti-tumor effects in an orthotopic mouse model. GL261-luciferase (GL261-Luc) cells were stereotactically implanted into the brains of Balb/C mice to establish intracranial tumors (Fig. 4A)^37,45^. To assess biodistribution and pharmacokinetics, EcomPC–IFN, native IFN-α, and the non-virus-like Encap cage were conjugated to Cy7 for near-infrared fluorescence tracking (Fig. 4A, B). Whole-body imaging revealed comparable systemic distribution across all groups, with prominent Cy7 fluorescence detected in the ventral view, particularly in the kidneys and heart, indicative of renal clearance (Fig. 4C, D). However, only EcomPC–IFN-treated mice exhibited a strong and sustained Cy7 signal in the brain, visible from 0.2 to 36 hours post-injection in dorsal views (Fig. 4C). In contrast, neither Cy7-labeled IFN-α nor Encap demonstrated detectable brain accumulation, underscoring their limited BBB permeability. Ex vivo imaging of major organs further confirmed these findings: while kidney fluorescence was consistently high across all treatment groups, only the EcomPC–IFN group showed substantial brain fluorescence (Fig. 4D). These results collectively highlight the ability of EcomPC to efficiently and durably transport therapeutic payloads across the BBB and into glioma tissue—overcoming a major barrier in central nervous system drug delivery and significantly extending the residence time of IFN-α within the tumor microenvironment.

**Fig 4.**
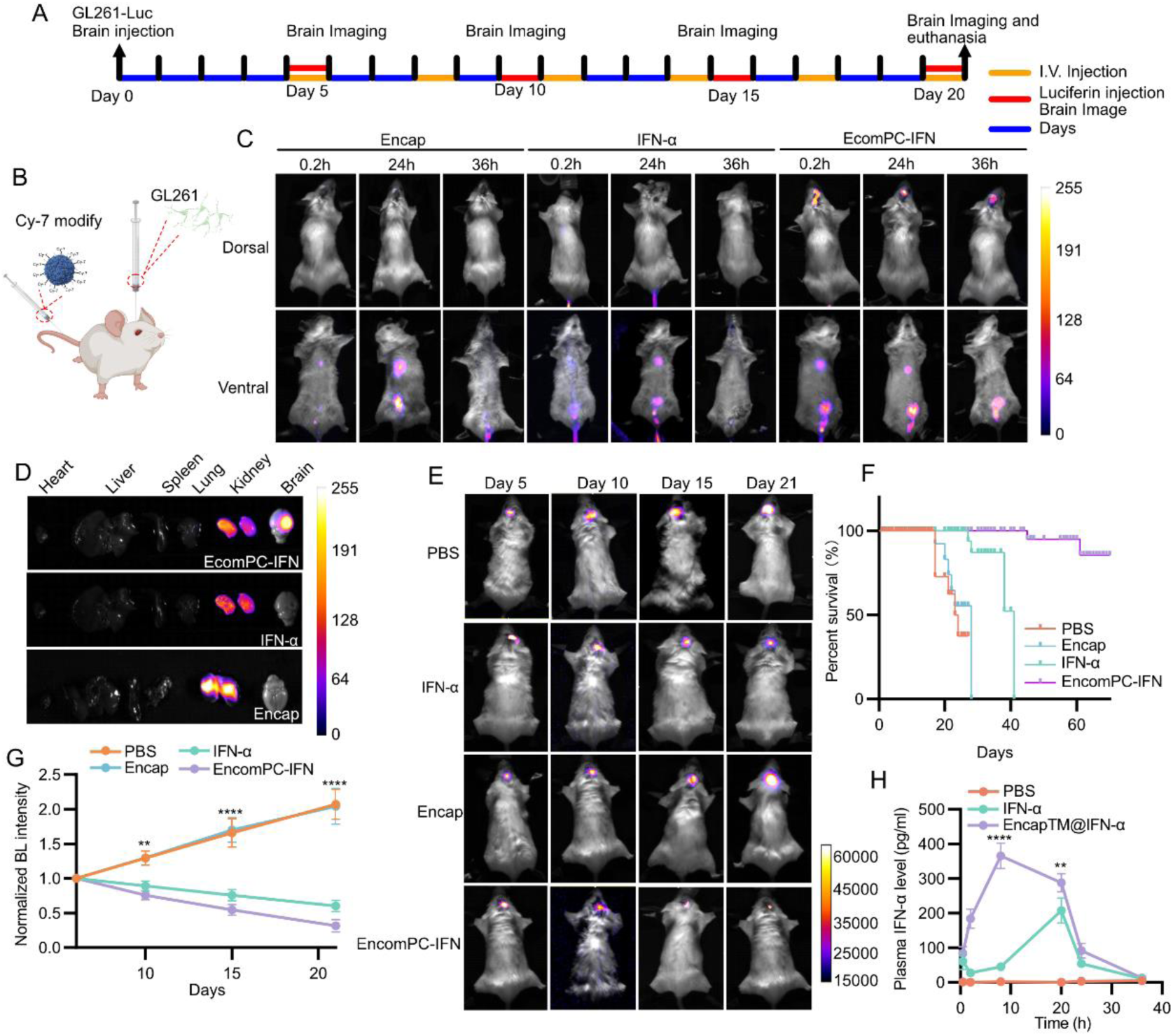
EcomPC-IFN exhibits enhanced BBB permeability, tumor targeting, and anti-GBM efficacy in vivo. A. Schematic timeline of the in vivo experimental protocol. GL261-Luc cells were stereotactically implanted to establish an orthotopic GBM mouse model. Mice received intravenous injections of recombinant protein constructs every other day. Bioluminescence imaging was performed on days 5, 10, 15, and 20 post-tumor implantations. B. Illustration of the biodistribution study. Recombinant proteins were conjugated with Cy7 fluorophore and administered via tail vein injection. C. Representative whole-body fluorescence images showing biodistribution of Cy7-labeled Encap, native IFN-α, and EcomPC-IFN at 0.2, 24-, and 36-hours post-injection. Images were captured from both dorsal and ventral views (n = 6 mice per group). D. Ex vivo fluorescence imaging of major organs 36 hours post-injection, confirming biodistribution patterns (n = 6). E. Bioluminescence imaging of tumor progression in GBM-bearing mice treated with PBS, Encap, IFN-α, or EcomPC-IFN at the indicated time points (n = 6). F. Kaplan–Meier survival curves of GBM-bearing mice following treatment. Mice were euthanized upon reaching humane endpoints as defined by ethical guidelines (n = 10). G. Quantification of normalized bioluminescence intensity, indicating tumor burden over time (n = 6). H. Serum IFN-α concentrations measured by ELISA at 1, 8-, 20-, 24-, and 36-hours post-injection in healthy mice treated with PBS, native IFN-α, or EcomPC-IFN (n = 3 per group).

Further therapeutic efficacy of EcomPC-IFN in vivo was assessed. Intracranial tumors beared Mice received intravenous injections of EcomPC-IFN, native IFN-α, blank Encap, or PBS every 3 days (Fig. 4A). Tumor progression was monitored by bioluminescence intensity (BLI) following intraperitoneal injection of D-luciferin on days 5, 10, 15, and 21 post-tumor implantations (Fig. 4A). BLI revealed a progressive increase in tumor signal intensity in both PBS- and blank Encap-treated groups^44,46^, with approximately a 2-fold increase in radiance from day 5 to day 20, indicating rapid tumor growth in the absence of effective treatment. In the group treated with native IFN-α, BLI levels remained largely unchanged over the 20-day period, suggesting modest tumor growth suppression but no significant regression (Fig. 4E, G). In contrast, mice treated with EcomPC-IFN exhibited a marked and sustained reduction in tumor bioluminescence, with signal intensity decreasing by approximately 80% relative to baseline (day 5), indicating substantial tumor regression (Fig. 4E, G). Body weight was monitored throughout the study as an indicator of general health. Tumor-bearing mice in control groups experienced >10% weight loss with disease progression, whereas EcomPC- IFN treated mice showed progressive weight gain, suggesting reduced tumor burden (Fig. S13). Furthermore, overall survival was evaluated in glioblastoma-bearing mice following the aforementioned treatment regimens. Kaplan–Meier survival analysis revealed the median survival time increased from 22 days in the PBS group and 40 days in the native IFN-α group to over 60 days in the EcomPC-IFN-treated group (Fig. 4F). These results demonstrate that systemic administration of EcomPC-IFN not only effectively penetrates the BBB but also elicits potent anti-tumor effects in vivo, surpassing the therapeutic efficacy of native IFN-α in this orthotopic GBM model.

As anticipated, intravenous administration of EcomPC–IFN resulted in significantly higher serum IFN-α levels compared to native IFN-α and PBS controls, peaking at ∼380 pg/mL at 8 hours post-injection (Fig. 4H). Sustained systemic IFN-α release has been shown to enhance antitumor synergy in cancer therapy^47^. Notably, elevated cytokine levels persisted for up to 36 hours in the EcomPC–IFN group, whereas native IFN-α rapidly declined to near-baseline levels (Fig. 4H). These results suggest that EcomPC not only enables efficient brain-targeted delivery but also prolongs systemic IFN-α exposure, consistent with previous findings that VLPs can stimulate systemic IFN-α responses^48^. Importantly, the prolonged cytokine circulation and viral-mimetic architecture raised concerns regarding systemic toxicity. However, histological examination of major organs, including the heart, liver, spleen, lungs, and kidneys, revealed no pathological abnormalities (Fig. S14), and hemolysis assays confirmed the absence of erythrocyte damage (Fig. S12). Collectively, these findings demonstrate that EcomPC-IFN combines extended systemic cytokine activity with excellent biocompatibility and in vivo safety.

### Targeted Delivery of IFN-α via EcomPC Nanocages Promotes GBM Apoptosis

To further assess the histopathological impact of EcomPC–IFN, we performed post- mortem analysis of brain tissues collected at the endpoint of our experimental timeline (Fig. 4A). BLI was decreased after EcomPC-IFN treatment, following whole-brain H&E revealed a substantial reduction in tumor signal following EcomPC–IFN treatment, which was corroborated by H&E staining of whole-brain sections. These analyses showed marked decreases in both tumor burden and invasive spread in the EcomPC–IFN group (Fig. 5A). Notably, tumor cells at the invasive margins displayed condensed and fragmented nuclei, morphological hallmarks of apoptosis, alongside enhanced cytoplasmic eosinophilia. These apoptotic features were largely absent or less pronounced in the PBS, Encap, and native IFN- α groups. (Fig. 5A).

**Fig 5.**
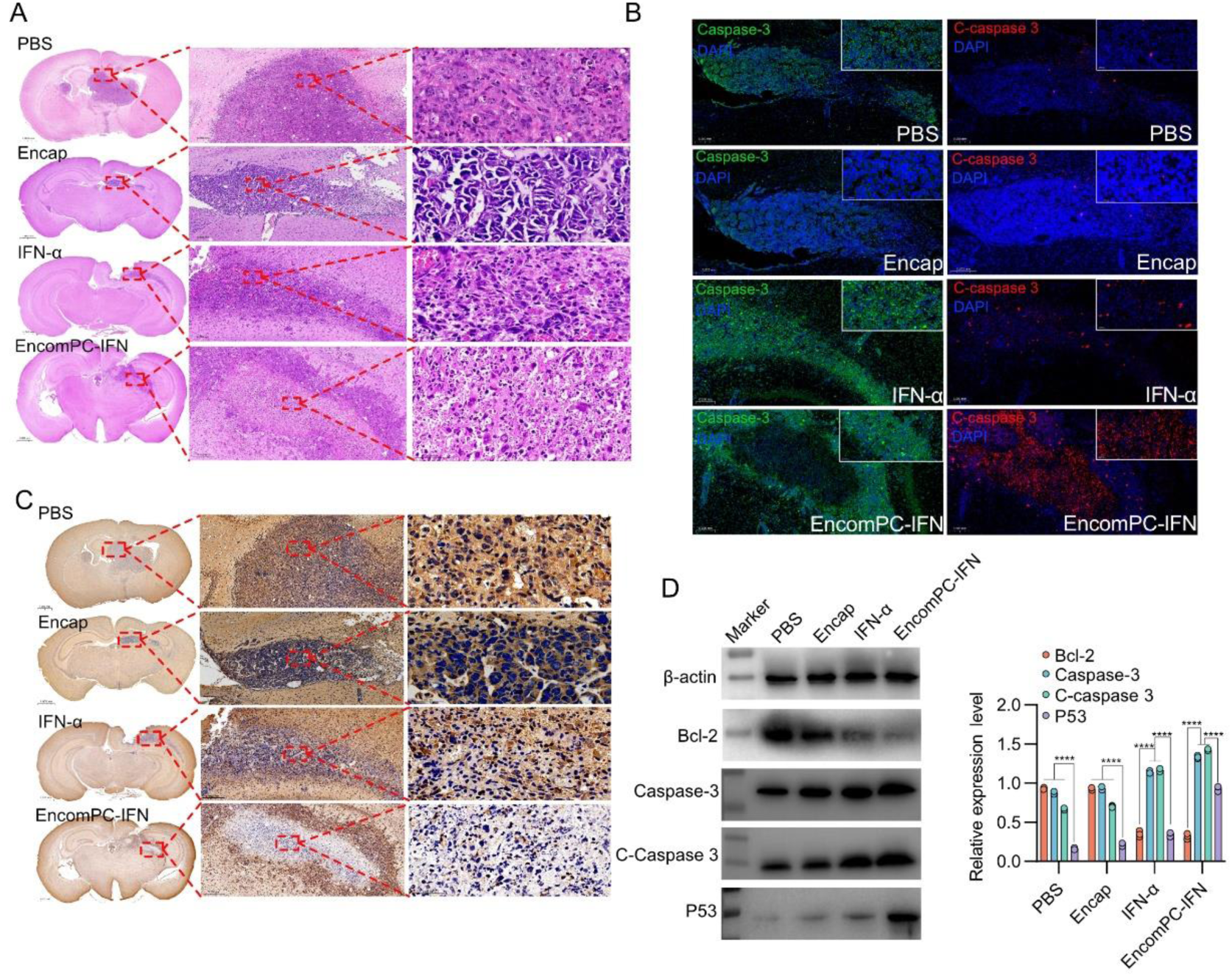
Targeted Delivery of EcomPC-IFN Induces Apoptosis in vivo. A. Representative H&E staining of whole-brain sections from glioblastoma-bearing mice treated with PBS, Encap, native IFN-α, or EcomPC-IFN. B. Immunohistochemical staining of total caspase-3 and cleaved caspase-3 in tumor regions. While total caspase-3 was broadly expressed across all groups, cleaved caspase-3 was significantly upregulated in the tumor core of EcomPC-IFN treated mice, indicating apoptotic execution. C. Immunohistochemical staining of BCL-2 reveals strong anti-apoptotic expression in PBS, Encap, and native IFN-α groups, but markedly reduced levels in tumors from EcomPC-IFN treated mice. D. Western blot analysis and quantification of whole-brain lysates shows elevated expression of p53, total caspase-3, and cleaved caspase-3, alongside suppressed BCL-2 expression following treatment. (n = 3; mean ±SEM; p < 0.01, one-way ANOVA).

As a clinically approved cytokine, IFN-α is known to induce tumor cell apoptosis through the Fas–Caspase–p53–BCL-2 signaling axis^49^. Consistent with this mechanism, immunohistochemical staining showed that total caspase-3 was present across all treatment groups; however, cleaved caspase-3, a key marker of apoptotic execution, was markedly upregulated within tumor cores following EcomPC–IFN administration (Fig. 5B, D). In contrast, minimal activation was observed in control groups. Furthermore, BCL-2 expression was significantly downregulated in EcomPC–IFN–treated tumors, whereas high levels were retained in PBS, Encap, and native IFN-α groups (Fig. 5C). This inverse relationship between cleaved caspase-3 and BCL-2 suggests that EcomPC–IFN promotes apoptosis by both activating pro-apoptotic signaling and suppressing anti-apoptotic defenses^45^. To validate these findings at the molecular level, we conducted Western blot analysis of whole-brain lysates. EcomPC–IFN treatment led to substantial upregulation of total and cleaved caspase-3, along with increased p53 expression and suppressed BCL-2 levels, consistent with immunohistochemical data (Fig. 5C and D). Collectively, these results provide mechanistic evidence that EcomPC-mediated IFN-α delivery activates the caspase–p53–BCL-2 apoptotic pathway, thereby enhancing tumor cell death in vivo.

Inflammatory responses are critical biological parameters to consider following VLP administration, especially given that coronavirus-mimicking structures may elicit innate immune activation^50^, and have been associated with neuroinflammatory events, including encephalitis, in certain coronaviruses such as OC43^18^. To evaluate the potential neuroinflammatory effects of EcomPC–IFN, we analyzed the expression levels of key inflammatory markers in whole-brain tissues. Interestingly, EcomPC–IFN treatment led to a significant upregulation of TNF-α, whereas both NF-κB and IL-6 levels were notably downregulated (Fig. S15). This pattern suggests that while EcomPC–IFN triggers localized immune activation, it does not induce a broad pro-inflammatory cascade. The selective increase in TNF-α, a cytokine involved in anti-tumor immunity, may contribute to the observed immunomodulatory and therapeutic effects, whereas the suppression of IL-6 and NF-κB implies a reduced risk of systemic or chronic neuroinflammation. These findings support the immune-regulating rather than immune-overactivating nature of EcomPC–IFN, highlighting its safety profile for CNS-targeted therapies.

### Targeted Delivery of IFN-α of EcomPC remodeling immune landscape of GBM tumor site

GBM establishes a profoundly immunosuppressive microenvironment characterized by the accumulation of regulatory T cells, infiltration of myeloid-derived suppressor cells, and the secretion of inhibitory cytokines, all of which collectively impair innate and adaptive immune responses and facilitate immune evasion by tumor cells^1,9^. This environment significantly impairs both innate and adaptive immune responses, allowing tumor cells to evade immune surveillance^1,9^. In this study, we designed EcomPC to deliver IFN-α directly to the tumor site, leveraging the dual immunomodulatory properties of both the cytokine and the viral-like architecture. IFN-α is well known to promote CD8⁺ T cell recruitment and cytotoxic activation^40,42^, while virus-mimicking nanostructures can engage pattern recognition receptors (PRRs) on innate immune cells, potentially triggering immune activation^23^. We therefore investigated whether EcomPC-mediated IFN-α delivery could reprogram the GBM immune milieu beyond inducing tumor cell apoptosis. Given the established role of local immunosuppressive mechanisms in GBM, including microglia-mediated cytokine secretion, impaired antigen presentation, and suppression of T cell activity, we first examine the Iba-1⁺ microglia and CD45⁺ leukocyte in tumor site^51^. Our results showed that treatment with EcomPC-IFN led to a marked increase in intratumoral CD45⁺ leukocyte infiltration, accompanied by a significant reduction in Iba-1⁺ microglial density (Fig. 6A). These changes were markedly more pronounced than those induced by native IFN-α or non-functionalized encapsulin controls (Fig. 6A and Fig. S16), suggesting that the coronavirus-mimicking scaffold synergizes with IFN-α to overcome local immunosuppression. Together, these results indicate that EcomPC–IFN not only delivers therapeutic cytokines across the BBB but also reprograms the GBM immune microenvironment toward a more immunostimulatory state.

**Fig 6.**
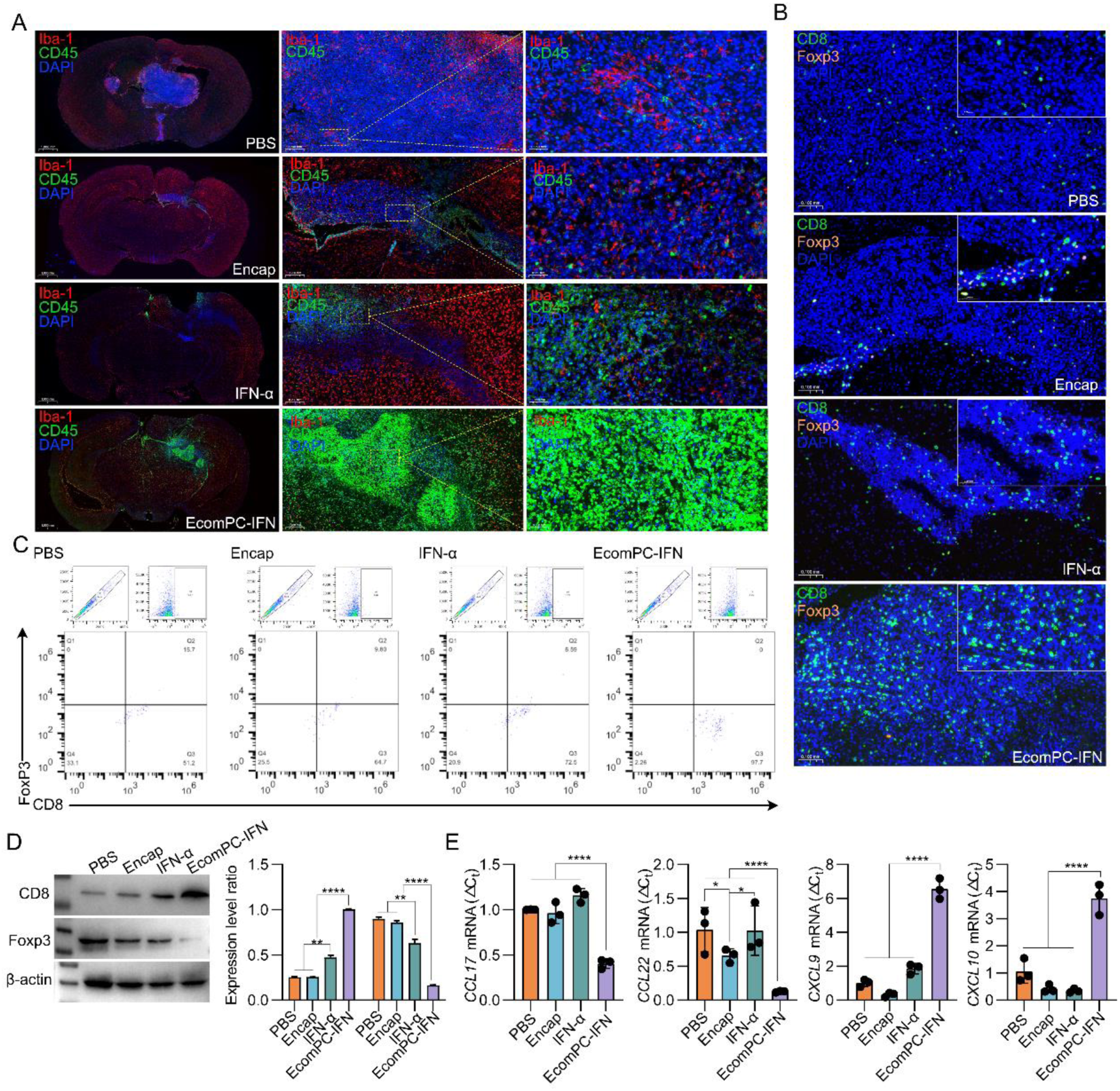
Targeted delivery of IFN-α via EcomPC nanocages reshapes the immune landscape of the glioma microenvironment. A. Whole-brain immunofluorescence staining for Iba-1 (microglia, red) and CD45 (leukocytes, green) revealed limited microglial infiltration across all groups. ECoMPC–IFN treatment significantly increased intratumoral CD45⁺ immune cell accumulation while further reducing microglial density. B. Dual immunofluorescence staining of CD8 and Foxp3 within tumor regions demonstrated a marked increase in CD8⁺ T cell infiltration and a concurrent reduction of Foxp3⁺ Tregs following EcomPC-IFN treatment, resulting in a significantly elevated CD8⁺/Foxp3⁺ ratio compared to controls. C. Flow cytometry analysis of brain-infiltrating CD45⁺ leukocytes confirmed the expansion of CD45⁺CD8⁺ cytotoxic T cells and reduction of CD45⁺Foxp3⁺ Tregs in the EcomPC–IFN group. D. Western blot analysis of whole-brain lysates showed increased CD8 expression and decreased FOXP3 levels following EcomPC–IFN treatment, corroborating histological and cytometric data. E. Quantitative PCR analysis of immune-regulatory chemokines revealed significant upregulation of *CXCL9* and *CXCL10*, alongside reduced expression of *CCL17* and *CCL22*, in EcomPC–IFN treated brains, indicating enhanced recruitment of effector T cells and suppression of Treg-attracting signals.

Elevated numbers of CD45⁺ tumor-infiltrating leukocytes are commonly observed in GBM, yet this infiltrate is skewed toward immunosuppressive phenotypes that blunt effective anti-tumor immunity^9,15^. To delineate how EcomPC-IFN reshapes this compartment, we next quantified the relative abundance of Foxp3⁺ regulatory T cells and CD8⁺ cytotoxic T lymphocytes within the CD45⁺ population. EcomPC-IFN treatment substantially increased CD8⁺ T cell infiltration while markedly decreasing Foxp3⁺ Treg presence, resulting in a significantly elevated intratumoral CD8⁺/ Foxp3⁺ ratio (Fig. 6B, C). Native IFN-α induced only modest changes in this ratio, and minimal immune infiltration was observed in PBS, Encap and other control groups (Fig. 6B, C and Fig. S16). Western blot analysis of whole-brain lysates further supported these results, revealing increased CD8 protein expression and decreased Foxp3 levels in EcomPC-IFN treated mice (Fig. 6D).

To further delineate the molecular underpinnings of the immune reprogramming observed following EcomPC-IFN treatment, we performed qPCR analysis on tumor-bearing brain tissues to assess the expression of key chemokines involved in T cell recruitment and immune regulation. Notably, the expression of *CXCL9* and *CXCL10*, both interferon-inducible chemokines critical for the recruitment of CXCR3⁺ effector T cells, was significantly upregulated in the EcomPC-IFN group compared to all controls, including native IFN-α. Conversely, the mRNA levels of *CCL17* and *CCL22*, chemokines known to mediate the recruitment of immunosuppressive Foxp3⁺ regulatory T cells, were markedly reduced following EcomPC-IFN administration (Fig. 6E and Fig. S17). These transcriptional changes further support the immunofluorescence and flow cytometry data, suggesting that EcomPC- mediated delivery of IFN-α not only enhances cytotoxic T cell infiltration but also suppresses Treg-attracting chemokine expression, thereby reshaping the tumor microenvironment toward a more immunostimulatory and anti-tumor phenotype.

## Discussion

Despite decades of investigation, the clinical translation of IFN-α for GBM remains hindered by poor brain penetration, rapid systemic clearance, and dose-limiting toxicity^8,14,52^. Addressing these limitations requires not only improved pharmacokinetic profiles^41^ but also platforms that preserve biological activity while targeting the complex architecture of the central nervous system^52^. In this work, we introduce EcomPC, a virus-mimicking, protein-based nanocage system, as a rationally engineered solution that integrates targeted delivery, molecular stability, and immunological precision.

Inspired by the structural tropism of neurotropic viruses, such as SARS-CoV-2^22^ and Zika virus^17^, EcomPC leverages key design elements found in viral systems, namely, symmetric nanogeometry and multivalent surface presentation, to mimic their ability to traverse biological barriers^16,22^. Unlike conventional nanocarriers, which often rely on passive diffusion or non-specific endocytosis, the viral-like architecture of EcomPC is hypothesized to engage active transcytotic pathways via size- and shape-dependent vesicular transport. This is supported by our observation that only fully assembled nanocages bearing both spike-like surface features^16,19^ and glioma-targeting peptides achieve efficient translocation across the BBB and accumulation within orthotopic GBM tissue^6^. Control constructs lacking either the architectural scaffold or targeting ligands fail to recapitulate this behavior, underscoring the requirement of cooperative structural and biochemical cues for BBB penetration.

Beyond physical delivery, EcomPC enables intracellular release and functional persistence of protein therapeutics. By engineering the encapsulin scaffold with a flexible linker and cysteine-to-serine substitutions^31^, we enhanced structural plasticity and minimized disulfide-mediated aggregation, thereby expanding the cargo accommodation capacity without compromising cage stability. Although the upper size limit for encapsulation remains to be precisely delineated, our DLS and TEM data indicate successful encapsulation of protein cargos up to ∼20–25 kDa (eGFP is 27 kDa, considering partial encapsulation). This suggests that EcomPC possesses sufficient internal cavity volume (∼24–30 nm diameter) and pore accessibility to accommodate small-to-medium sized proteins in monomeric or moderately folded states^31,34^.

Importantly, this work reveals that spatially restricted IFN-α delivery via EcomPC induces not only potent direct cytotoxicity but also a systems-level immunological reprogramming of the tumor microenvironment. Sustained local IFN-α exposure activates canonical p53- mediated apoptotic pathways in tumor cells, consistent with the known pro-apoptotic role of type I interferons^47,53^. Concurrently, we observed a shift in the immune landscape characterized by increased infiltration of cytotoxic CD8⁺ T cells and a marked suppression of immunosuppressive Tregs, indicating a reversal of the tolerogenic niche that typically defines glioma^40,42^. Although virus-like protein cages have been shown in some contexts to engage pattern recognition receptors (PRRs)^21,54^, we did not observe overt immune activation from the EcomPC scaffold alone. Thus, the immunological effects observed here are most parsimoniously explained by the sustained, spatially controlled action of IFN-α-a cytokine well known to augment antigen presentation, promote effector T cell activation, and inhibit tumor-induced immune tolerance^40^.

The therapeutic significance of these findings extends beyond conventional cytokine delivery. IFN-α is a pleiotropic immunomodulator that promotes dendritic cell maturation, increases antigen cross-presentation, and boosts natural killer cell cytotoxicity, all of which converge to amplify tumor antigen visibility and immune effector recruitment^40,42^. However, its systemic application has historically been hampered by short half-life, peripheral toxicity, and immune cell exhaustion^41^. EcomPC overcomes these barriers by localizing IFN-α bioactivity within the tumor site, thereby minimizing peripheral cytokine storm risk while sustaining therapeutically relevant exposure for immune education. Notably, the observed remodeling of the tumor microenvironment was not accompanied by evidence of overt innate immune activation from the EcomPC scaffold itself. This suggests that EcomPC, despite its viral mimicry, acts as an “immunological enabler” rather than a danger signal, delivering cytokines in a temporally and spatially resolved manner that favors coordinated immune activation rather than indiscriminate immune stimulation.

In conclusion, EcomPC represents more than a delivery vehicle-it is a structurally informed, immunologically active nanoplatform that bridges protein engineering with translational immunotherapy. By mimicking evolutionary principles of viral architecture, this system overcomes longstanding barriers in CNS drug delivery, and opens new avenues for combining apoptotic and immune pathways in a spatially and temporally controlled manner. Future work will expand this paradigm to multi-cargo strategies, combination immunotherapies, and broader applications in neuro-oncology and beyond.

## Materials and methods

### Chemical agents

The Encap sequence and primers were synthesized by Sangon Biotech Co., Ltd. The plasmids were synthesized within the laboratory. The ligase, restriction enzymes, and polymerase used were all purchased from Thermo Scientific. The β-actin antibody (Lot: AC22073004), Bcl-2 antibody (Lot: AC230918049), phosphorylated P53 antibody (Lot: AC231016067), caspase3 antibody (Lot: GB115600), and cleaved caspase3 (Lot: GB11532) were obtained from Servicebio. Additionally, the HRP-conjugated goat anti-rabbit IgG (Lot: AC2401261007), Alexa Fluor 488-conjugated goat anti-rabbit IgG (Lot: GB25303), CY3- conjugated goat anti-rabbit IgG (Lot: GB21303), bovine serum albumin (BSA, Lot: GC305010), normal rabbit serum (concentrated, Lot: G1209), and DAPI staining reagent (Lot: G1012) were also purchased from Servicebio. The CCK- 8 assay kit and Live/Dead staining kit were provided by Dalian Meilun Biotechnology Co., Ltd. The 7-AAD cell viability detection reagent was obtained from Beyotime Biotechnology. The CD45 antibody (Lot: bsm- 30251A-PE-cy5), CD8a antibody (Lot: bsm-30396A-PE), and Foxp3 (Lot: bsm-30294M- FITC) antibody were all purchased from Beijing Obsen Biotechnology Co., Ltd.

### Molecular Cloning

To construct the EcomPC system, we designed and synthesized a series of expression plasmids. The Encap_mut1_-Int^C^-ST, Encap_mut2_-Int^C^-ST, and Encap-Int^C^-ST gene fragments were PCR-amplified and inserted into the pBAD vector (Invitrogen), which confers ampicillin resistance and allows arabinose-inducible expression in *E. coli*^55^. These constructs were designed to include a SpyTag (ST) at the C-terminus of Int^C^-Encap-ST for downstream modular ligation^27^.

Separately, coding sequences for the POI, and the targeting peptides LinTT1-SC and LinTT1-mut-SC, were amplified and inserted into the pET-28a vector under control of the T7 promoter. These sequences were engineered to include an Int^N^ domain to enable intracellular protein trans-splicing with the encapsulin subunits.

All recombinant constructs were confirmed by restriction enzyme digestion and Sanger sequencing to verify correct insertion, reading frame, and sequence fidelity. High-quality plasmid DNA was prepared using an established protocol for subsequent transformation into *E. coli* BL21(DE3) competent cells.

### Protein expression and purification

For co-expression of encapsulin scaffolds and POI cargos, *E. coli* BL21(DE3) cells co-transformed with pBAD–Int^C^–Encap–ST and pET-28a–POI–Int^N^ plasmids were cultured in Luria-Bertani (LB) medium supplemented with ampicillin (50 mg/mL) and kanamycin (50 mg/mL) at 37 °C until reaching an optical density at 600 nm (OD₆₀₀) of 0.5–0.6. Expression of the POI was induced by adding 0.5 mM isopropyl β-D-1-thiogalactopyranoside (IPTG), followed by incubation at 18 °C for 5 hours^56,57^. Subsequently, encapsulin expression was induced by the addition of 0.1% (w/v) L-arabinose, and cultures were further incubated overnight at 18 °C to allow in vivo self-assembly. For single-plasmid expression of other recombinant proteins (LinTT1–SpyCatcher, mutant LinTT1–SC), cells harboring the corresponding pET-28a constructs were grown under similar conditions, with IPTG induction (0.5 mM) at OD₆₀₀ = 0.5–0.6, followed by overnight incubation at 18 °C. Bacterial cells were harvested by centrifugation at 8,000 × g for 20 minutes at 4 °C, and the resulting pellets were resuspended in lysis buffer (50 mM NaH₂PO₄, 300 mM NaCl, 10 mM imidazole, pH 8.0). Cells were lysed by ultrasonication on ice, and lysates were cleared by centrifugation (12,000 × g, 30 minutes, 4 °C)^58^. The supernatants were subjected to nickel-nitrilotriacetic acid (Ni-NTA) affinity chromatography under native conditions. Proteins were eluted using imidazole- gradient elution buffer (typically 250 mM imidazole, pH 8.0). Protein purity and molecular weight were assessed by SDS–PAGE followed by Coomassie Brilliant Blue staining, and target identity was confirmed by Western blot using an anti-His-tag antibody. All proteins were stored at 4 °C in phosphate-buffered saline (PBS; 50 mM Na₂HPO₄, 100 mM NaCl, pH 7.4) prior to use^56,59^.

### EcomPC preparation

To generate the final EcomPC nanostructures, equimolar solutions of POI contented encap protein solution (1.5 mg/mL) and TP-SC (1.5 mg/mL) were mixed at varying volume ratios (10:1, 8:2, 5:5, 4:6, 2:8, and 1:10) to evaluate conjugation efficiency via SpyTag/SpyCatcher coupling. Mixtures were incubated overnight at 16 °C to ensure complete reaction. The extent of conjugation was assessed by SDS–PAGE followed by Coomassie Brilliant Blue staining. Among the tested ratios, a 2:8 volumetric ratio of POI contented encap protein to ST-SC yielded the most efficient conjugation, characterized by near-complete consumption of unreacted ST-SC and the appearance of distinct higher-molecular-weight bands. These optimized conditions were subsequently used for downstream applications, including cellular uptake, targeting specificity, and therapeutic efficacy assays.

### Characterizations of EcomPC Nanostructures

The morphology of EcomPC constructs (EcomPC-eGFP, EcomPC–IFN-α, Encap, etc.) was examined using TEM. Samples (10 μL) were applied onto carbon-coated copper grids (Electron Microscopy Sciences) and incubated for 1 min at room temperature. Excess liquid was blotted off using filter paper. The grids were then negatively stained with 5 μL of 1% (w/v) uranyl acetate for 1 min, followed by blotting and air-drying overnight. Imaging was performed using a JEOL JEM-1400 transmission electron microscope (JEOL Ltd.) operated at an accelerating voltage of 120 kV. Additional imaging was conducted using a Tecnai 10 TEM (FEI) equipped with a LaB₆ cathode (100 kV) and an UltraScan 1000 CCD camera (Gatan) for high-resolution acquisition.

The hydrodynamic diameter and size distribution of EcomPC nanoparticles were measured by DLS using a Zetasizer Nano ZS90 system (Malvern Instruments). Measurements were conducted at 25 °C with a fixed scattering angle of 90° and a laser wavelength of 633 nm. Prior to analysis, protein samples were filtered through a 0.22 μm pore-size membrane (Millipore) to remove particulates. Data were processed using Zetasizer software, and all measurements were performed in triplicate to ensure reproducibility.

### Cell Culture

NIH-3T3, HaCaT, GL261, and HeLa cell lines were obtained from commercial sources and used for in vitro experiments^37^. Cells were maintained in Dulbecco’s Modified Eagle Medium (DMEM; Gibco, USA) supplemented with 10% fetal bovine serum (FBS; Gibco, USA) and 1% penicillin–streptomycin (HyClone, SV30010, USA). All cultures were incubated at 37 °C in a humidified atmosphere containing 5% CO₂.

### Cytotoxicity and Cell Viability Assays

To assess cytotoxicity, GL261 and HeLa cells were seeded in 96-well plates at a density of 5 ×10³cells per well and allowed to adhere overnight. For EcomPC-eGFP evaluation, GL261 cells were treated with eGFP-LinTT1, Encap, or EcomPC-eGFP at concentrations of 2.5, 5, 10, or 20 μM for 24 h. For EcomPC-IFN evaluation, GL261 and HeLa cells were treated with free IFN-α or EcomPC-IFN at identical concentrations and timepoints. After treatment, 10% (v/v) Cell Counting Kit-8 (CCK-8, MeilunBio) solution was added to each well and incubated for 1–2 h at 37 °C. Absorbance at 450 nm was measured using a microplate reader (BioTek), and cell viability was normalized to the PBS-treated control group.

Cell viability was further assessed using a calcein-AM/propidium iodide (PI) dual-staining assay (MeilunBio). Following treatment under the same conditions as the CCK-8 assay, cells were incubated with calcein-AM (2 μM) and PI (2 μM) for 5 min at 37 °C. Fluorescence images were acquired using an inverted fluorescence microscope (Nikon Eclipse Ti2), and viable (green) vs. dead (red) cell populations were quantified.

For the scratch wound migration assay, GL261 and HeLa cells were seeded in 6-well plates and grown to ∼80% confluence. A linear scratch was introduced using a sterile 20-μL pipette tip (defined as time 0 h), and detached cells were removed by gentle PBS washing. Cells were then treated with IFN-α or EcomPC-IFN at 10 or 20 μM and cultured for up to 24 h. Wound closure was imaged at 0, 12, and 24 h using phase-contrast microscopy. The wound area was quantified using ImageJ software, and migration was expressed as a percentage of initial wound area.

### Blood–Brain Barrier (BBB) Transwell Penetration Assay

To evaluate the translocation efficiency of nanoformulations across a model BBB, a cellular Transwell co-culture system was established using HaCaT and GL261 cells^37^. MilliCell Hanging Cell Culture Inserts (0.4 μm pore size; Millipore, USA) were placed in 24- well plates to serve as the upper compartment. HaCaT cells (5 × 10⁴ cells/well) were seeded in the upper chamber with 200 μL of DMEM supplemented with 10% fetal bovine serum (FBS), while the lower chamber was filled with 600 μL of the same medium to maintain osmotic balance^37^. GL261 cells (5 × 10⁴ cells/well) were cultured on sterile circular glass coverslips placed at the bottom of the lower chamber to simulate the brain parenchymal interface. After 24 h of culture, the medium was replaced with serum-free DMEM in both compartments. EcomPC-eGFP or other groups was added to the upper chamber at final concentrations of 2.5, 5, or 10 μM (based on protein concentration of 1.5 mg/mL). The co-culture was incubated at 37 °C with 5% CO₂ for an additional 24 h to allow for potential transcytosis. Following incubation, cells in the lower chamber were stained with propidium iodide (PI, 2 μM) and DAPI to assess viability and visualize nuclei. Fluorescence microscopy was used to detect intracellular GFP signal in GL261 cells, indicating successful traversal of EComPC-eGFP across the in vitro BBB model^37^.

### Flow Cytometry Analysis of Cytotoxicity

To assess the cytotoxic effects of EcomPC–IFN-α on glioma cells, GL261 cells were seeded into 6-well plates and treated with either free IFN-α or EcomPC–IFN-α at final concentrations of 10 μM or 20 μM. After 24 h of incubation at 37 °C, cells were harvested, washed twice with cold PBS, and resuspended in 1× binding buffer. Cell viability was assessed using 7-Aminoactinomycin D (7-AAD, Beyotime, China) staining according to the manufacturer’s instructions. After 10 min of incubation at 37 °C in the dark, samples were immediately analyzed using a BD SORP ARIA III flow cytometer (BD Biosciences). A minimum of 10,000 events per sample were collected. Data acquisition and analysis were performed using FlowJo software (TreeStar Inc.), and the percentage of 7-AAD⁺ cells was used as an indicator of compromised membrane integrity and cell death.

### Haemolysis Assay

To evaluate the hemocompatibility of the coronavirus-mimicking protein nanocages, a hemolysis assay was performed using whole blood collected from healthy Balb/c mice. Fresh blood was diluted to a final concentration of 5% (v/v) in PBS. Test samples were prepared at 20% (w/v) in PBS. Triton X-100 (0.1%) and PBS served as the positive and negative controls, respectively. Equal volumes (100 μL) of diluted blood and test samples were mixed in 1.5 mL microcentrifuge tubes and incubated at 37 °C for 60 minutes in a thermostatic metal bath. After incubation, the mixtures were centrifuged at 1,000 ×g for 12 minutes. The supernatants (100 μL) were transferred to a 96-well plate, and absorbance at 540 nm was measured using a microplate reader to quantify released hemoglobin^56,57^.

### Intracranial Mouse Glioma Model

Male BALB/c mice (20–25 g) were housed under standard conditions and acclimatized for one week prior to experimentation. To establish an orthotopic glioma model, animals were anesthetized using an isoflurane vaporizer system (RWD, R510IP) and secured in a stereotactic frame. After scalp depilation and antiseptic preparation, a midline incision was made to expose the cranial surface. A glioma cell suspension was prepared by resuspending 5 × 10⁵ GL261-acGFP-Fluc cells in 20 μL of 50% Matrigel. Cells were stereotactically injected into the right striatum using the following coordinates relative to Bregma: +0.98 mm anterior, +1.5 mm lateral to midline, and −2.5 mm ventral from the skull surface^10^. The injection needle was positioned at a slight angle toward the coronal suture to minimize reflux. Following implantation, the burr hole was sealed with sterile bone wax (Fine Science Tools, Germany), and the scalp was closed using 6-0 sutures (Chenghe Micro Instrument Factory, China). Mice were monitored until full recovery and housed under pathogen-free conditions for subsequent in vivo experiments.

### Bioluminescent (Luciferase) In Vivo Imaging in Small Animals

On postoperative day 5, tumor engraftment was verified by bioluminescence imaging following intravenous injection of D-luciferin potassium salt (15 mg/mL). Mice were anesthetized with 2% isoflurane in 100% oxygen at a flow rate of 1 L/min. Imaging was conducted 10 minutes post-injection of D-luciferin (75 mg/kg) using the IVIS Spectrum small animal imaging system (BG-gdsAUTO730, BayGene Biotech Co., Ltd)^60^.

### In Vivo Imaging Targeting Assay of Cy7-Labeled Protein

Following successful establishment of the glioma mouse models, protein nanocomplexes (IFN-α, EcomPC-IFN and native Encap) were each labeled with Cy7 dye. Specifically, 1 mL of each protein solution was mixed with 100 μL of coupling reagent and 0.5 mg of Cy7 dye (excitation: 745 nm; emission: 800 nm), then incubated in the dark for 30 minutes^61^. The reaction mixture was transferred to an ultrafiltration tube and centrifuged at 10,000 rpm for 10 minutes to remove unbound dye, after which the retentate was resuspended to 1 mL with PBS. Glioma-bearing mice were randomly assigned to control and experimental groups. The control group received protein nanocomplexes lacking targeting peptides, while the experimental group received LinTT1-targeted complexes. Each mouse was administered 100 μL of the respective formulation via tail vein injection, designated as time zero (0 h). Fluorescence imaging was performed using 740 nm excitation at 0.2 h, 24 h, and 36 h post-injection to monitor biodistribution. Upon completion of imaging, mice were euthanized, and major organs were harvested for ex vivo fluorescence imaging to assess tissue-specific drug targeting.

### Serum IFN-α Concentration Measurement

Plasma concentrations of IFN-α were quantified by ELISA. Mice were administered IFN-α or EcomPC-IFN intravenously via tail vein at a dose of 150 μg/kg. Blood samples (100 μL each) were collected from the heart under isoflurane anesthesia at predetermined time points (0.5, 2, 8-, 20-, 24-, and 36-hours post-injection). Samples were allowed to clot at 4°C for 30 minutes, followed by centrifugation at 100 × g for 15 minutes to isolate plasma, which was then stored at –80°C until analysis. Baseline levels were established using plasma from untreated mice and subtracted from all experimental values. Quantification was performed using a human IFN-α2 ELISA kit (PBL Interferon Source) according to the manufacturer’s instructions. Pharmacokinetic analysis was conducted using GraphPad Prism version 9.0.0.

### Western Blot Analysis

Total proteins were extracted from mouse brain tissues using a protein extraction kit (Solarbio, BC3710-100T, China), and protein concentrations were determined using a BCA assay. Equal amounts of protein were separated by sodium dodecyl sulfate-polyacrylamide gel electrophoresis (SDS-PAGE) and transferred onto nitrocellulose membranes. Membranes were blocked with 5% non-fat milk or BSA in TBST to prevent nonspecific binding, followed by overnight incubation at 4°C with primary antibodies targeting β-actin, Bcl-2, p53, caspase-3, cleaved caspase-3, CD8a, Foxp3, NF-κB, and IL-6. After washing, membranes were incubated with HRP-conjugated secondary antibodies (anti-rabbit or anti-mouse, EnVision system) for 1 hour at room temperature. Protein bands were visualized using enhanced chemiluminescence (ECL) reagents and imaged with a digital imaging system. Band intensities were quantified using ImageJ software, and protein expression levels were normalized to β-actin as a loading control.

### RNA Extraction and Quantitative Real-Time PCR

Total RNA was isolated from mouse brain tissues using the RNA Easy Fast Tissue Total RNA Extraction Kit (TIANGEN) according to the manufacturer’s protocol. Reverse transcription was carried out using the FastKing gDNA Dispelling RT SuperMix (TIANGEN), which includes a genomic DNA removal step to ensure specificity. Quantitative real-time PCR (qRT-PCR) was performed using FastReal qPCR PreMix (SYBR Green) (TIANGEN) on a CFX96 Real-Time PCR Detection System (Bio-Rad). Primer sequences used for amplification are provided in Supplementary Table 2. Gene expression levels were normalized to GAPDH, and relative fold changes were calculated.

### Flow Cytometric Analysis of Tumor-Infiltrating Immune Cells

Brain tumor tissues were harvested and mechanically dissociated using a syringe plunger to generate a single-cell suspension. The suspension was transferred to a 15 mL conical tube and centrifuged at 1,500 rpm for 5 min at 4 °C. The supernatant was discarded, and red blood cells were lysed by resuspending the pellet in 5 mL of RBC lysis buffer, followed by incubation at room temperature for 8 min. The lysed suspension was passed through a 70 μm cell strainer to remove debris. Cells were then centrifuged again at 1,500 rpm for 10 min and resuspended in PBS. For surface marker staining, aliquots of the cell suspension were incubated with anti-CD45 and anti-CD8a antibodies at 4 °C for 30 min in the dark. For intracellular FoxP3 staining, cells were first fixed in 4% paraformaldehyde at room temperature, washed, and permeabilized in PBS containing 0.5% BSA. After centrifugation at 1,200 rpm for 5 min, cells were incubated with the anti-Foxp3 antibody under permeabilizing conditions. All stained cells were finally resuspended in 500 μL PBS and analyzed using a CytoFLEX LX flow cytometer (Beckman Coulter). Data acquisition and analysis were performed using FlowJo software (Tree Star Inc., Ashland, OR).

### Histological Analysis and H&E Staining

On day 21 post-treatment, mice were euthanized by cervical dislocation and perfused with PBS. Brain, heart, liver, spleen, lungs, and kidneys were rapidly excised and fixed in 4% paraformaldehyde (PFA) at 4 °C for 24–48 h. Fixed tissues were dehydrated through a graded ethanol series (70%, 80%, 90%, 95%, and 100%), cleared in xylene, and embedded in paraffin according to standard protocols. Paraffin-embedded tissues were sectioned at 4–5 μm using a rotary microtome (Leica RM2235), deparaffinized in xylene, and rehydrated through descending ethanol concentrations. Sections were stained with hematoxylin (5 min) for nuclear visualization and eosin (2 min) for cytoplasmic contrast. After dehydration and xylene clearing, slides were mounted using neutral balsam and coverslipped. Images were acquired using a Leica DMi8S inverted microscope equipped with a high-resolution CCD camera.

### Immunofluorescence and Immunohistochemistry Staining

Paraffin-embedded tissue sections (4–5 μm) were deparaffinized in xylene (2 × 10 min) and rehydrated through a descending ethanol gradient (100%, 95%, 70%) followed by distilled water. Antigen retrieval was performed by heating slides in either citrate buffer (pH 6.0) or Tris-EDTA buffer (pH 9.0) in a microwave oven (95–100 °C, 15–20 min), followed by cooling to room temperature. For intracellular antigen detection, sections were permeabilized with 0.1% Triton X-100 and blocked with 5% normal rabbit serum for 1 h at room temperature. Sections were then incubated overnight at 4 °C in a humidified chamber with the following primary antibodies diluted in blocking buffer: anti-Caspase-3, anti-Cleaved Caspase-3, anti-CD45, anti-CD8a, and anti-FoxP3. After three PBS washes (5 min each), slides were incubated for 1 h at room temperature with species-specific Alexa Fluor-conjugated secondary antibodies (488/594/647, 1:500; protected from light). Nuclear counterstaining was performed using DAPI (1 μg/mL, 5 min). Sections were mounted with antifade medium and stored at 4 °C in the dark. Images were captured using a Leica DMi8S fluorescence microscope with identical acquisition settings across experimental groups.

For chromogenic detection, paraffin sections were processed similarly through deparaffinization, rehydration, antigen retrieval, and blocking. Sections were incubated overnight at 4 °C with rabbit monoclonal anti-Bcl-2 antibody diluted in 3% BSA/PBS. After washing (3 × 5 min, PBS), sections were incubated with HRP-conjugated anti-rabbit secondary antibodies (EnVision™ system, Dako) for 1 h at room temperature. Chromogenic development was performed using DAB substrate (5 min, RT), and the reaction was quenched with distilled water. Nuclei were counterstained with hematoxylin (30 sec), followed by dehydration in graded ethanol (70%, 95%, 100%), clearing in xylene, and mounting with resinous medium. Brightfield images were acquired using a Nikon Eclipse C1 microscope.

### Statistics and data reproducibility

All data are presented as mean ± standard error of the mean (SEM), unless otherwise indicated. Statistical analyses were performed using GraphPad Prism 10.1.2 and Origin 2021. For comparisons involving multiple groups, two-way analysis of variance (two-way ANOVA) was applied, followed by appropriate post hoc tests. Statistical significance was denoted as follows: *: p < 0.05, **: p < 0.01, ***: p < 0.001, ****: p < 0.0001. Image analysis and quantification were performed using Fiji (ImageJ) software. All experiments were independently repeated at least three times unless otherwise specified, and representative data are shown. No data points were excluded unless justified by experimental error or pre-established criteria.

## Acknowledgement

The animal experiments were approved by Inner Mongolia University (SYXK 2020-0006). This project was supported by a grant from National Natural Science Foundation (22365022); Inner Mongolia Natural Science Foundation (2022QN03014); Inner Mongolia Science and Technology Project (2022ZY0050); Inner Mongolia Youth Science and Technology Talent Support Project (NUYT23092); Science and Technology Leading Talent Team in Inner Mongolia Autonomous Region (2022LJRC0009); Inner Mongolia Autonomous Region Science and Technology Project (2023YFHH0010). All authors have approved the final version of this manuscript. We wish to thank the Electron Microscopy Centre of Inner Mongolia University for the microscopy and microanalysis of our specimens. Ethical approval for the in vivo breast cancer and glioma model construction experiments was granted by the Institutional Animal Care and Use Committee of Inner Mongolia University.

## Conflict of Interest

The authors declare no conflict of interest.

## Data Availability Statement

The data supporting the findings and all plasmids and proteins are available upon reasonable request from the corresponding author.

## Supporting Information

The Supporting Information is available free of charge at: Supplementary.

SDS-PAGE and WB for proteins. TEM images of mutated EcomPC. SDS-PAGE validation of Encap-IFN/eGFP with LinTT1-SC. Structural prediction LinTT1 targeting peptides. CCK8 assay of G261 cell of EcomPC-eGFP. Live/dead staining of GL261 cell for EcomPC-IFN and other groups. Cell migration of GL261 cell. CCK8 assay for Hela cell. Cell migration of Hela cell. Hemolysis assay. H&E staining of the heart, liver, spleen, lung and kidney. WB for inflammatory factors. Immunofluorescence staining after native EcomPC treatment. The quantitative PCR analysis of native EcomPC. Amino acid sequences of genes used in the article. The qPCR primers used in the article.

## Notes

### Competing Interest Statement

The authors have declared no competing interest.

