## Supplemental Table 1 for "Rationally Engineered Coronavirus-mimicking Protein Nanocage Platform for Glioblastoma Targeted Delivery and Immunity Regulation"

+

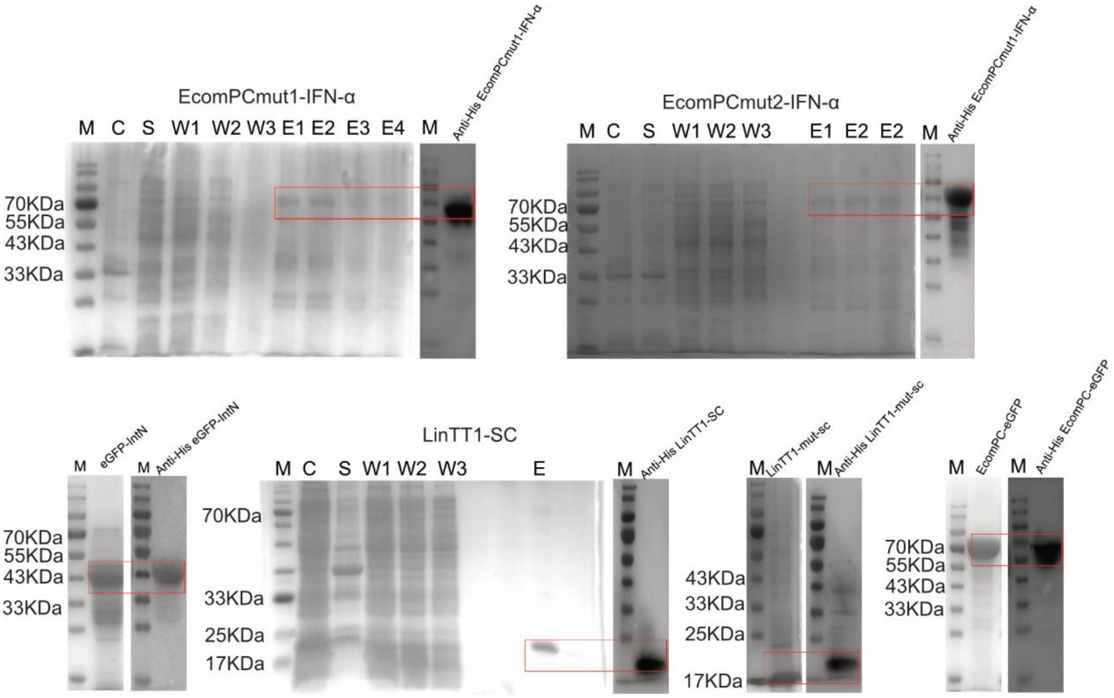


Figure. S1: Protein purification SDS-PAGE and WB validation of Encaps with mutation sites and proteins Encaps with eGFP、IFN-α conjugation.

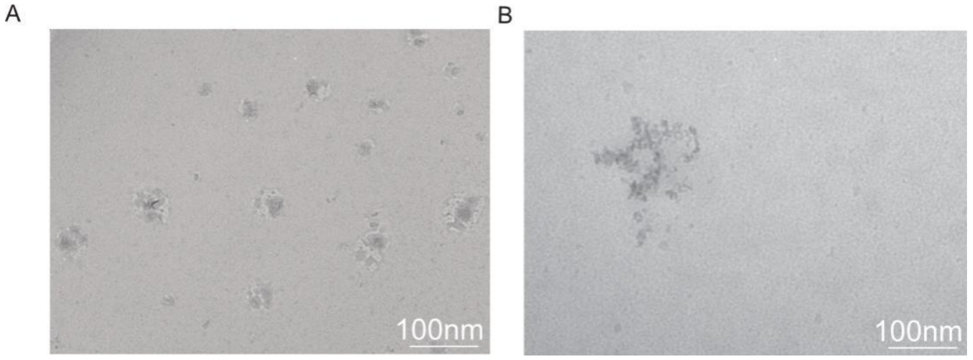

Figure. S2: Electron microscopy characterization of EcomPcmut1-IFN-α (A) and EcomPCmut2-IFN-α (B).

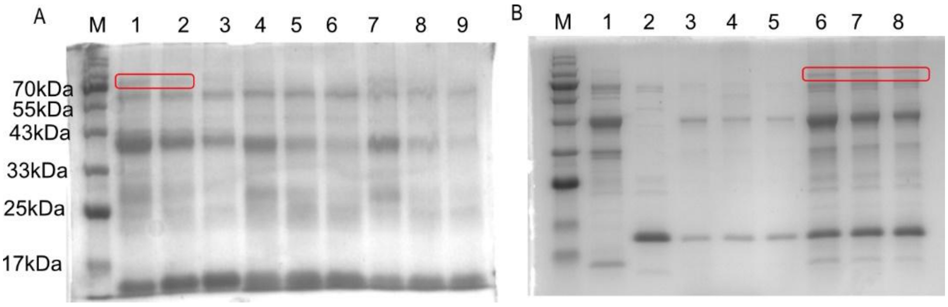


Figure. S3：Click Chemical Modification of Encap-IFN-α/eGFP with LinTTI-SC Recombinant Protein （A）Click Chemical Modification of Encap-eGFP with LinTTI-SC Recombinant Protein(mass ratio)：M ：Marker，1-2：the combination ratio of Encap-eGFP to LinTTI-SC at 2:8，3-4：the combination ratio of Encap-eGFP to LinTTI-SC at 8:2, 5：the combination ratio of Encap-eGFP to LinTTI-SC at 4:6, 6：the combination ratio of Encap-eGFP to LinTTI-SC at 5:5, 7：the combination ratio of Encap-eGFP to LinTTI-SC at 1:10, 8-9：the combination ratio of Encap-eGFP to LinTTI-SC at 10:1（B）Click Chemical Modification of Encap-IFN-α with LinTTI-SC Recombinant Protein(mass ratio)：M ：Marker，1：the combination ratio of Encap-IFN-α to LinTTI-SC at 4:6，2：the combination ratio of Encap-IFN-α to LinTTI-SC at 5:5, 3：the combination ratio of Encap-IFN-α to LinTTI-SC at 6:4, 4：the combination ratio of Encap-IFN-α to LinTTI-SC at 1:10, 5：the combination ratio of Encap-IFN-α to LinTTI-SC at 10:1, 6-8：the combination ratio of Encap-IFN-α to LinTTI-SC at 2:8.

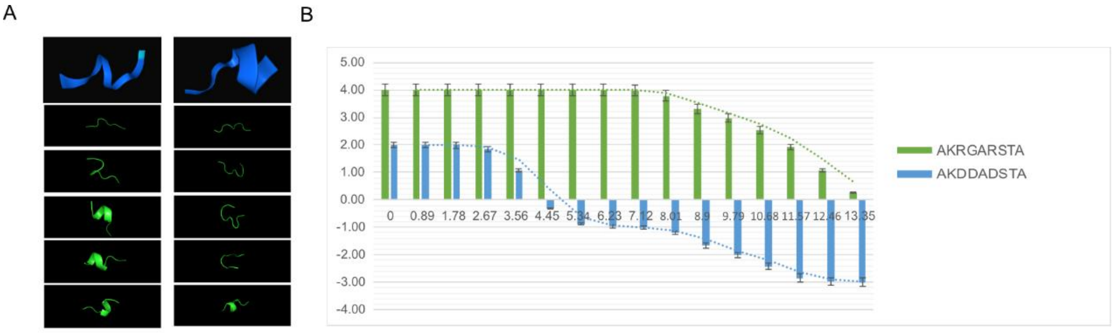


Figure. S4: Structural prediction (A)、 Electrostatic charge quantity and their changing trend (B) of LinTT1-mut-SC ( AKDDADSTA ) and LinT T1-SC (AKRGARSTA).

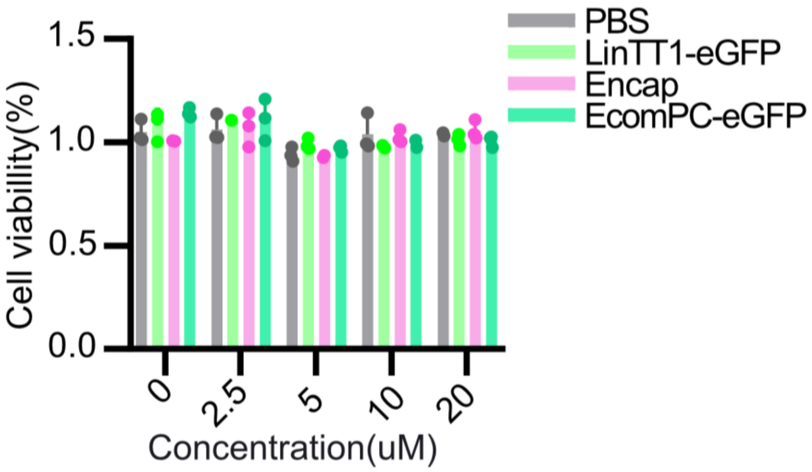


Figure. S5: The CCK8 assay of G261 cell by PBS, LinT T1, Encap, and EcomPC-eGFP at different concentration gradients.

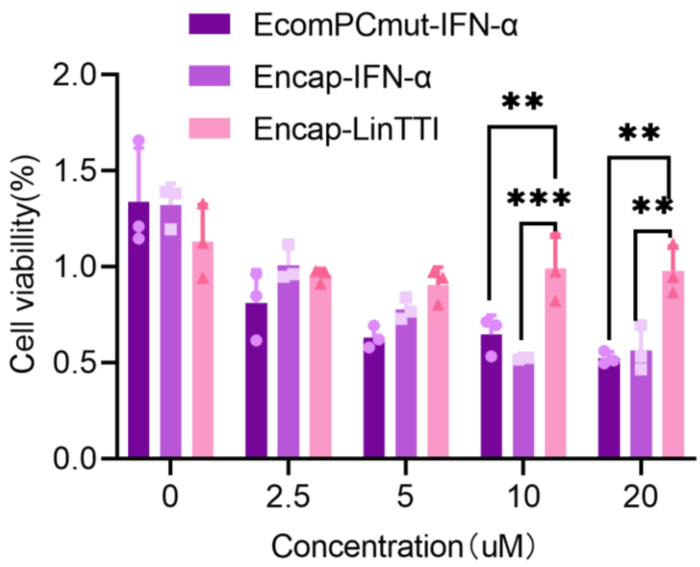


Figure. S6: The CCK8 assay of G261 cell by EcomPCmut-IFN-α、Encap-IFN-α and Encap-LinTTI at different concentration gradients.

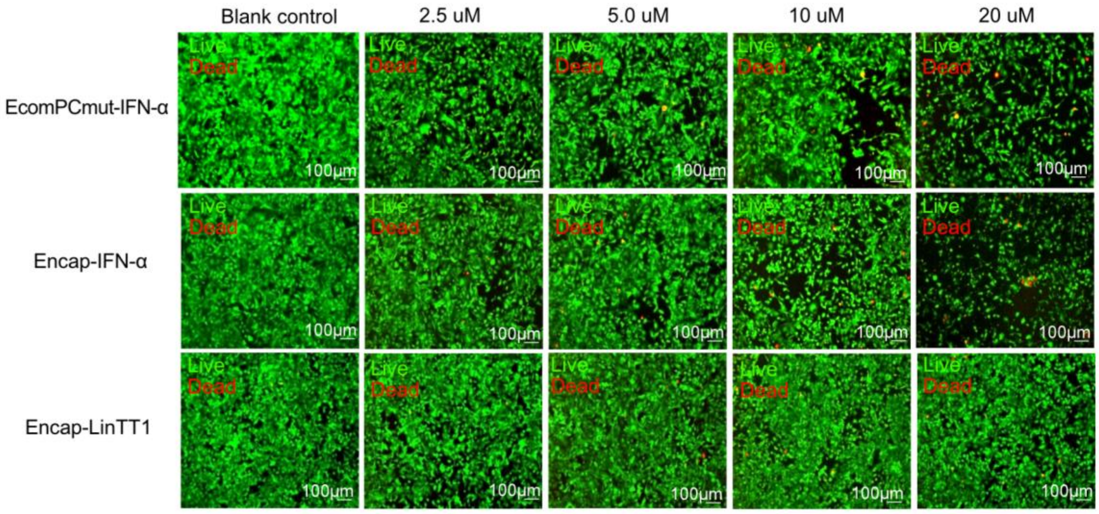


Figure. S7:Results of dead-viable staining of GL261 cells by EcomPCmut-IFN-α、Encap-IFN-α and Encap-LinTT1 at different concentration gradients.

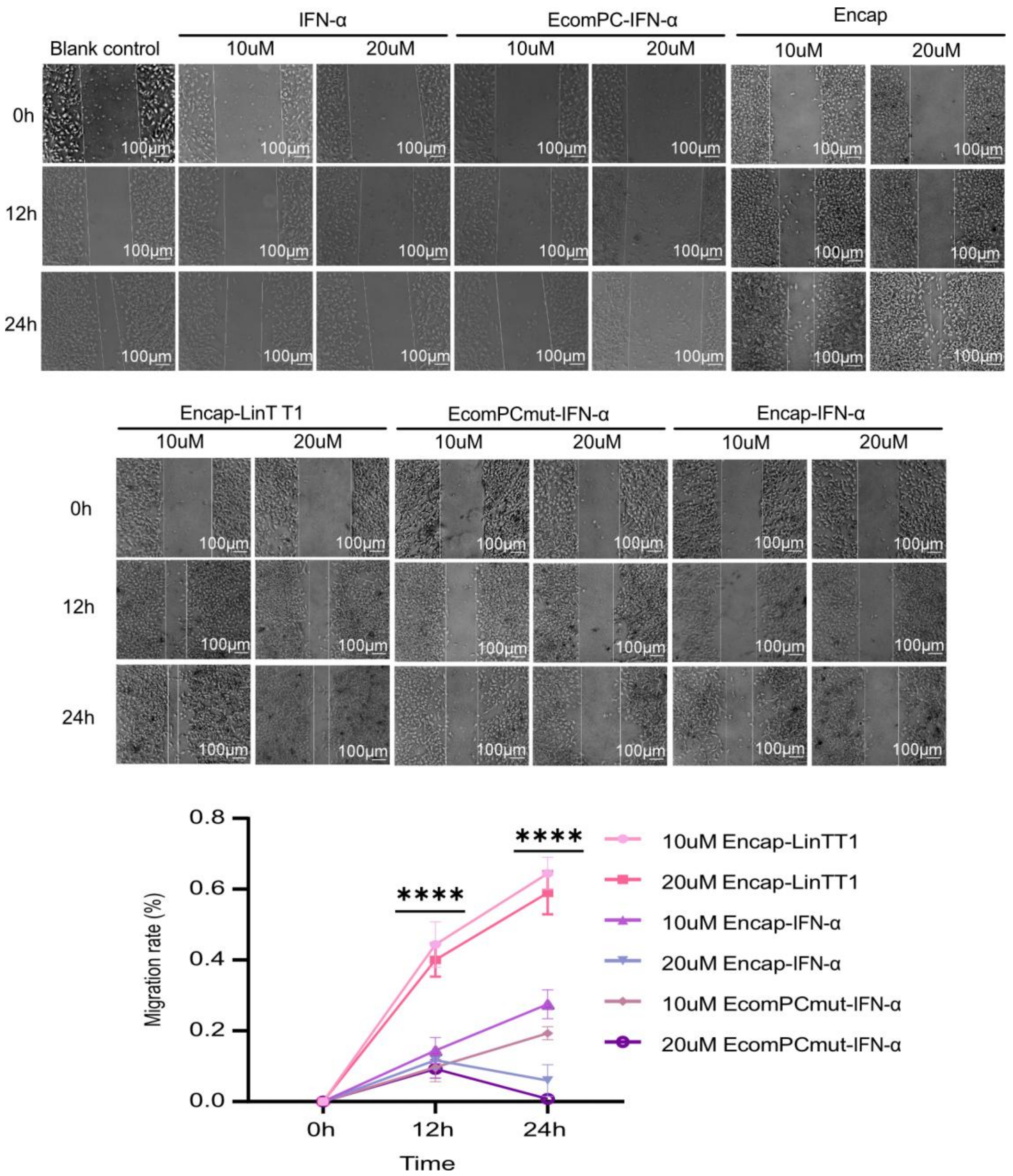


Figure. S8: The impact of IFN-α, EcomPC-IFN-α, Encap, Encap-LinTT1, EcomPCmut-IFN-α, Encap-IFN-α on G261 cell migration.

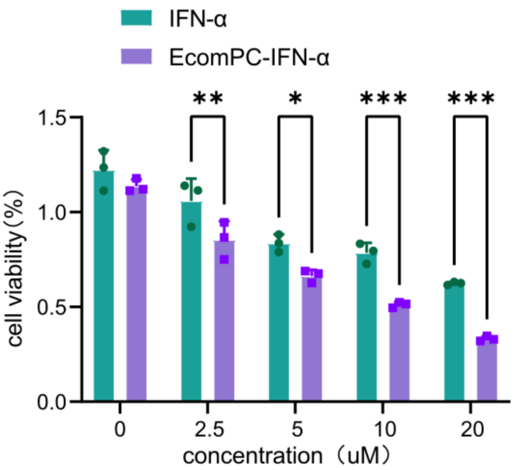


Figure. S9: The CCK8 assay of HeLa cell by IFN-α and EcomPC-IFN-α at different concentration gradients

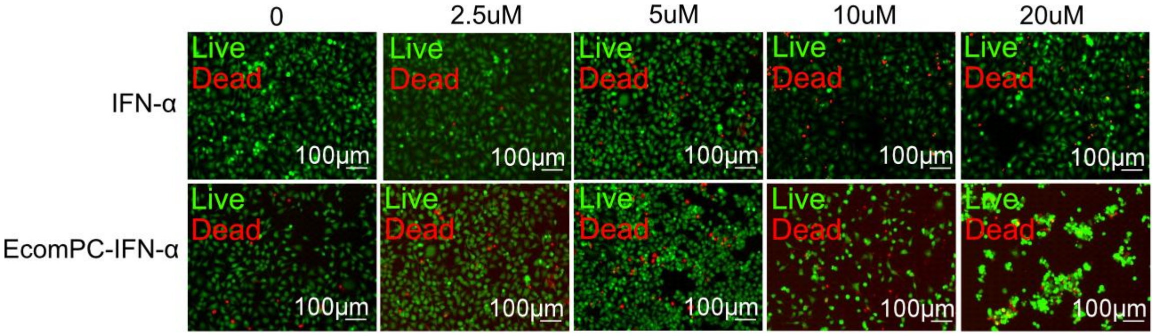


Figure. S10: Results of dead-viable staining of HeLa cells by IFN-α and EcomPC-IFN-α at different concentration gradients.

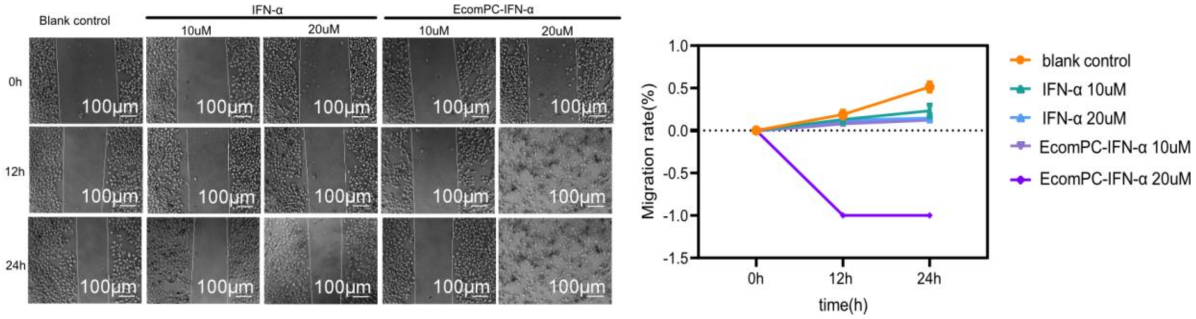


Figure. S11: The impact of PBS, Encap, IFN-α and EcomPC-IFN-α on HeLa cell migration.

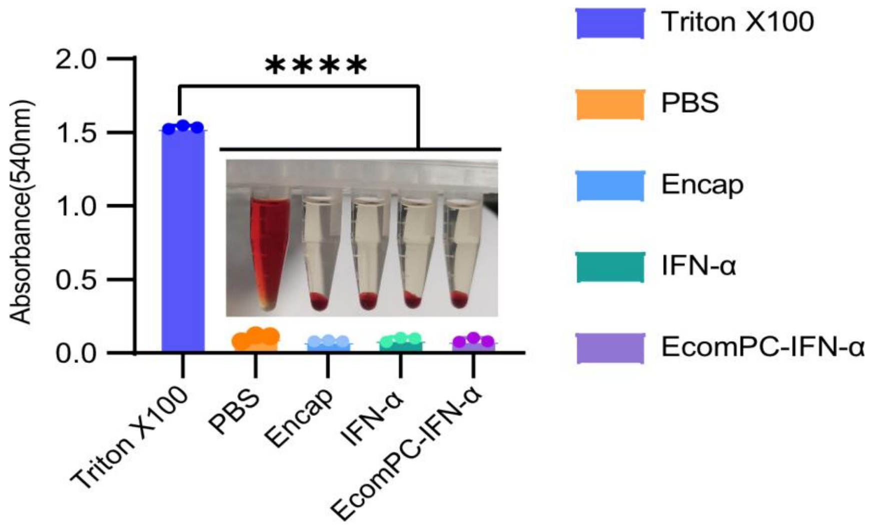


Figure. S12: Results of hemolysis assay for Triton-X100, PBS, Encap, IFN-α, and EcomPC-IFN-α.

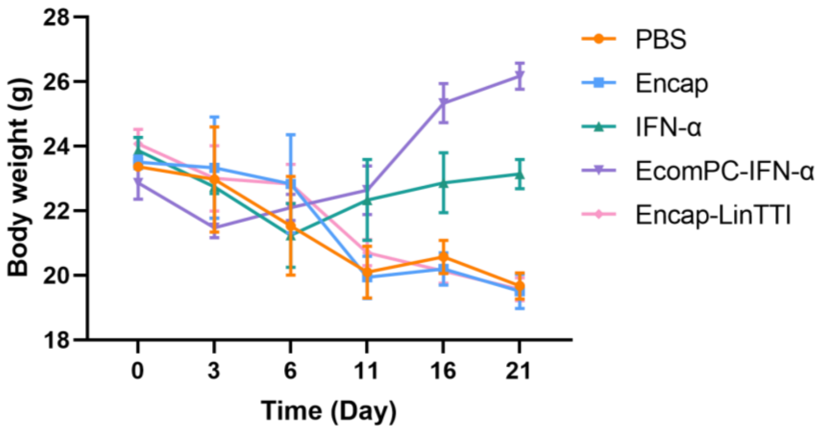


Figure. S13: Body weight curves of mice in each treatment group.

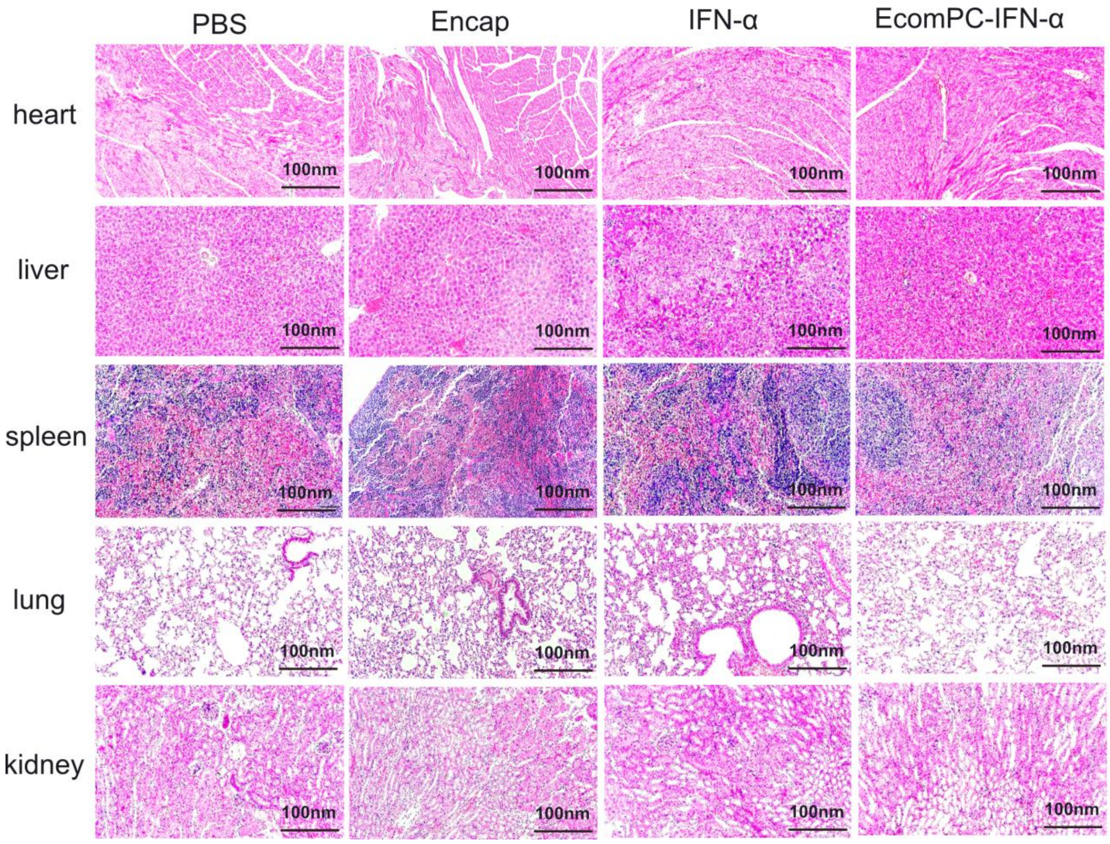


Figure. S14: HE staining of the heart, liver, spleen, lung and kidney of mice in each treatment group.

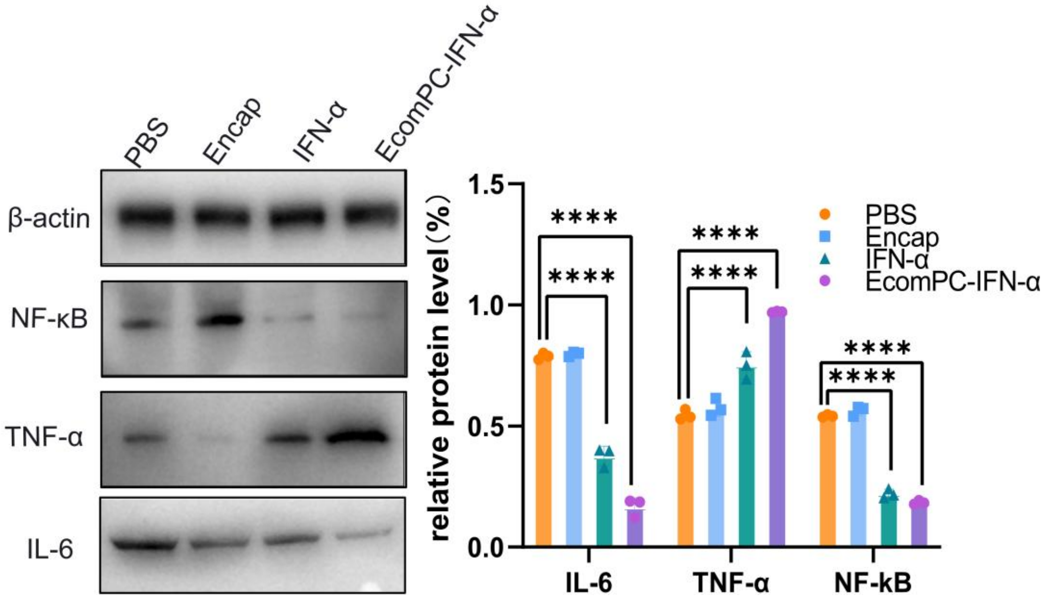


Figure. S15: WB validation of four groups of brain protein in mice with glioma tumors with PBS, Encap, IFN-α, EcomPC-IFN-α and their grayscale analysis.

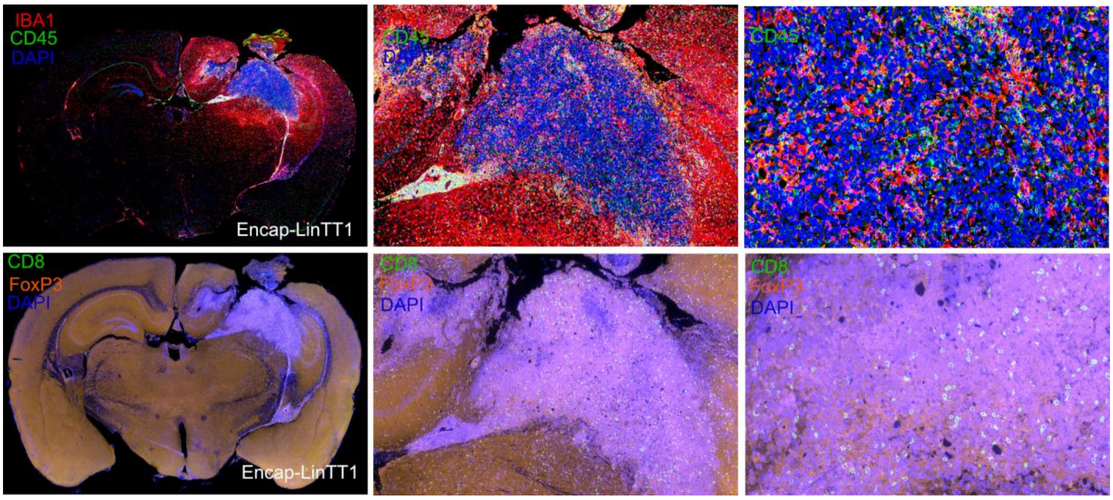


Figure. S16: Double immunofluorescence staining for Iba1 (microglia, red) and CD45 (leukocytes, green), CD8 and Foxp3 in Enca-LinTT1 treated whole brain sections.

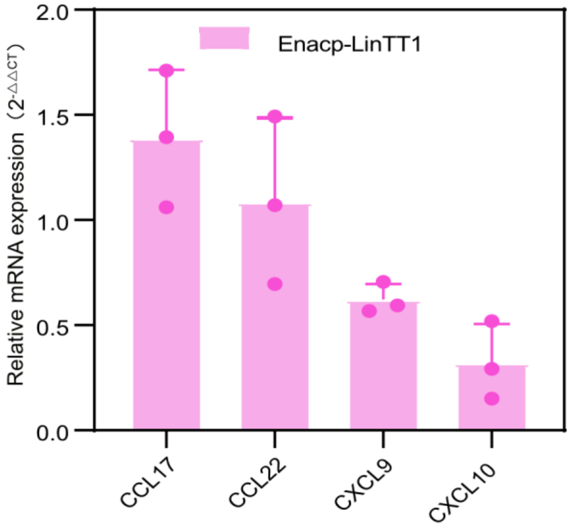


Figure. S17: The quantitative PCR analysis results of the immunoregulatory chemokines CXCL9, CXCL10, CCL17 and CCL22 in the brain treated with Encap-LinTT1.

| Name of gene(s) | **Amino acid sequences** |
|---|---|
| IFN-α-IntN | CDLPQTHSLGSRRTLMLLAQMRRISLFSCLKDRHDFGFPQEEFGNQ FQKAETIPVLHEMIQQIFNLFSTKDSSAAWDETLLDKFYTELYQQL NDLEACVIQGVGVTETPLMKEDSILAVRKYFQRITLYLKEKKYSPC AWEVVRAEIMRSFSLSTNLQESLRSKEGNSGGGLVAGGSGGGSGT SAEYCLSYETEILTVEYGLLPIGKIVEKRIECTVYSVDNNGNIYTQPV AQWHDRGEQEVFEYCLEDGSLIRATKDHKFMTVDGQMLPIDEIFE RELDLMRVDNLPN |
| eGFP-IntN | VSKGEELFTGVVPILVELDGDVNGHKFSVSGEGEGDATYGKLTLK FICTTGKLPVPWPTLVTTLTYGVQCFSRYPDHMKQHDFFKSAMPE |

| | |
|---|---|
| | GYVQERTIFFKDDGNYKTRAEVKFEGDTLVNRIELKGIDFKEDGNILGHKLEYNYNSHNVYIMADKQKNGIKVNFKIRHNIEDGSVQLADHYQQNTPIGDGPVLLPDNHYLSTQSALSKDPNEKRDHMVLLEFVTAAGITLGMDELYKGNSGGGLVAGGSGGGSGTSAEYCLSYETEILTVEYGLLPIGKIVEKRIECTVYSVDNNGNIYTQPVAQWHDRGEQEVFEYCLEDGSLIRATKDHKFMTVDGQMLPIDEIFERELDLMRVDNLPN |
| IntC-Encap-ST | MIKIATRKYLGKQNVYDIGVERDHNFALKNGFIASNCFNGGGGSGGGGSGGGGSGTGGSEFLKRSFAPLTEKQWQEIDNRAREIFKTQLYGRKFVDVEGPGGGGGHHHHHHASGGGGGYGWEYAAHPLGEVEVLSDENEVVKWGLRKSLPLIELRATFTLDLWELDNLERGKPNVDLSSLEETVRKVAEFEDEVIFRGCEKSGVKGLLSFEERKIESGSTPKDLLEAIVRALSIFSKDGIEGPYTLVINTDRWINFLKEEAGHYPLEKRVEESLRGGKIITTPRIEDALVVSERGGDFKLILGQDLSIGYEDREKDAVRLFITETFTFQVVNPEALILLKFEPPPPLPPPPEPPPPGSAHIVMVDAYKPTK |
| LinTT1-SC | MVDTLSGLSSEQGQSGDMTIEEDSATHIKFSKRDEDGKELAGATMELRDSSGKTISTWISDGQVKDFYLYPGKYTFVETAAPDGYEVATAITFTVNEQGQVTVNGKATKGDAHIEFAKRGARSTA |
| IntC-Encap mut1-ST | MIKIATRKYLGKQNVYDIGVERDHNFALKNGFIASNCFNGGGGSGGGGSGGGGSGTGGSEFLKRSFAPLTEKQWQEIDNRAREIFKTQLYGRKFVDVEGPGGGGGHHHHHHASGGGGGYGWEYAAHPLGEVEVLSDENEVVKWGLRKSLPLIELRATFTLDLWELDNLERGKPNVDLSSLEETVRKVAEWEDEVYWRGCEKSGVKGLLSFEERKIESGSTPKDLVEAYVRALSIFSKDGIEGPYTLVINTDRWINFLKEEAGHYPLEKRVEESLRGGKIITTPRIEDALVVSERGGDFKLILGQDLSIGYEDREKDAVRLFITETFTFQVVNPEALILLKFEPPPPLPPPPEPPPPGSAHIVMVDAYKPTK |
| IntC-Encap mut2-ST | MIKIATRKYLGKQNVYDIGVERDHNFALKNGFIASNCFNGGGGSGGGGSGGGGSGTGGSEFLKRSFAPLTEKQWQEIDNRAREIFKTQLYGRKFVDVEGPGGGGGHHHHHHASGGGGGYGWEYAAHPLGEVEVLSDENEVVKWGLRKSLPLIELRATFTLDLWELDNLERGKPNVDLSSLEETVRKVAEKEDEVHKRGCEKSGVKGLLSFEERKIESGSTPKDLREAHVRALSIFSKDGIEGPYTLVINTDRWINFLKEEAGHYPLEKRVEESLRGGKIITTPRIEDALVVSERGGDFKLILGQDLSIGYEDREKDAVRLFITETFTFQVVNPEALILLKFEPPPPLPPPPEPPPPGSAHIVMVDAYKPTK |
| LinTT1-mut-SC | MVDTLSGLSSEQGQSGDMTIEEDSATHIKFSKRDEDGKELAGATMELRDSSGKTISTWISDGQVKDFYLYPGKYTFVETAAPDGYEVATAITFTVNEQGQVTVNGKATKGDAHIEFAKDDADSTA |
| LinTT1-eGFP | MVDTLSGLSSEQGQSGDMTIEEDSATHIKFSKRDEDGKELAGATMELRDSSGKTISTWISDGQVKDFYLYPGKYTFVETAAPDGYEVATAI |

| | |
|---|---|
| | TFTVNEQGQVTVNGKATKGDAHIGGSGGVSKGEELFTGVVPILVE LDGDVNGHKFSVSGEGEGDATYGKLTLKFICTTGKLPVPWPTLVT TLTYGVQCFSRYPDHMKQHDFFKSAMPEGYVQERTIFFKDDGNYK TRAEVKFEGDTLVNRIELKGIDFKEDGNILGHKLEYNYNSHNVYIM ADKQKNGIKVNFKIRHNIEDGSVQLADHYQQNTPIGDGPVLLPDN HYLSTQSALSKDPNEKRDHMVLLEFVTAAGITLGMDELYK |

Table. S1: Amino acid sequences of genes used in the article.

| | |
|---|---|
| β-actin-F1 | AGATTACTGCCCTGGCTCCTAG |
| β-actin-R1 | CATCGTACTCCTGCTTGCTGAT |
| CCL17-F1 | AGGGATGCCATCGTGTTTCT |
| CCL17-R1 | AGGTCATGGCCTTGGGTTTT |
| CCL22-F1 | AACCTTCTTGCTCCTCTGGA |
| CCL22-R1 | AAGCCCTTTGTGGTCCCATA |
| CXCL9-F1 | AGGGATGCCATCGTGTTTCT |
| CXCL9-R1 | AGGGAGGTGGACAACGTTTT |
| CXCL10-F1 | CCACGTGTTGAGATCATTGCC |
| CXCL10-R1 | GAGGCTCTCTGCTGTCCATC |

Table. S2: The qPCR primers used in the article.
